# Ancient viral genomes reveal introduction of HBV and B19V into Mexico during the transatlantic slave trade

**DOI:** 10.1101/2020.06.05.137083

**Authors:** Axel A. Guzmán-Solís, Daniel Blanco-Melo, Viridiana Villa-Islas, Miriam J. Bravo-López, Marcela Sandoval-Velasco, Julie K. Wesp, Jorge A. Gómez-Valdés, María de la Luz Moreno-Cabrera, Alejandro Meraz-Moreno, Gabriela Solís-Pichardo, Peter Schaaf, Benjamin R. tenOever, María C. Ávila-Arcos

## Abstract

After the European colonization of the Americas there was a dramatic population collapse of the Indigenous inhabitants caused in part by the introduction of new pathogens. Although there is much speculation on the etiology of the Colonial epidemics, direct evidence for the presence of specific viruses during the Colonial era is lacking. To uncover the diversity of viral pathogens during this period, we designed an enrichment assay targeting ancient DNA (aDNA) from viruses of clinical importance and applied it on DNA extracts from individuals found in a Colonial hospital and a Colonial chapel (16^th^ c. – 18^th^ c.) where records suggest victims of epidemics were buried during important outbreaks in Mexico City. This allowed us to reconstruct three ancient human parvovirus B19 genomes, and one ancient human hepatitis B virus genome from distinct individuals. The viral genomes are similar to African strains, consistent with the inferred morphological and genetic African ancestry of the hosts as well as with the isotopic analysis of the human remains, suggesting an origin on the African continent. This study provides direct molecular evidence of ancient viruses being transported to the Americas during the transatlantic slave trade and their subsequent introduction to New Spain. Altogether, our observations enrich the discussion about the etiology of infectious diseases during the Colonial period in Mexico.

## INTRODUCTION

European colonization in the Americas resulted in a frequent genetic exchange mainly between Native American populations, Europeans, and Africans (Aguirre-Beltrán, 2005; Rotimi et al., 2016; Salas et al., 2004). Along with human migrations, numerous new species were introduced to the Americas including bacterial and viral pathogens, which played a major role in the dramatic population collapse that afflicted the immunologically-naïve Indigenous inhabitants (Acuña-Soto et al., 2004; Lindo et al., 2016). Among these pathogens, viral diseases, such as smallpox, measles and mumps have been proposed to be responsible for many of the devastating epidemics during the Colonial period (Acuña-Soto et al., 2004). Remarkably, the pathogen(s) responsible for the deadliest epidemics reported in New Spain (the Spanish viceroyalty that corresponds to Mexico, Central America, and the current US southwest states) remains unknown and is thought to have caused millions of deaths during the 16^th^ century^4^. Indigenous populations were drastically affected by these mysterious epidemics, generically referred to as *Cocoliztli* (“pest” in Nahuatl)^6^, followed by Africans and to a lesser extent European people (Acuña-Soto et al., 2004; Malvido & Viesca, 1982; Somolinos d’Árdois, 1982). Symptoms of the 1576 *Cocoliztli* epidemic were described in autopsy reports of victims treated at the “Hospital Real de San José de los Naturales” (HSJN) (Malvido & Viesca, 1982; Wesp, 2017), the first hospital in Mexico dedicated specifically to treat the Indigenous population (Malvido & Viesca, 1982; Wesp, 2017) (Figure 1a-b). The symptoms described included high fever, severe headache, neurological disorders, internal and external bleeding, hepatitis and intense jaundice (Acuña-Soto et al., 2004; Malvido & Viesca, 1982; Somolinos d’Árdois, 1982). This has led some scholars to postulate that the etiological agent of the *Cocoliztli* epidemic was a hemorrhagic fever virus (Acuña-Soto et al., 2004; Marr & Kiracofe, 2000), although others have suggested that the symptoms could be explained by bacterial infections (Malvido & Viesca, 1982; Vågene et al., 2018).

**Figure 1.**
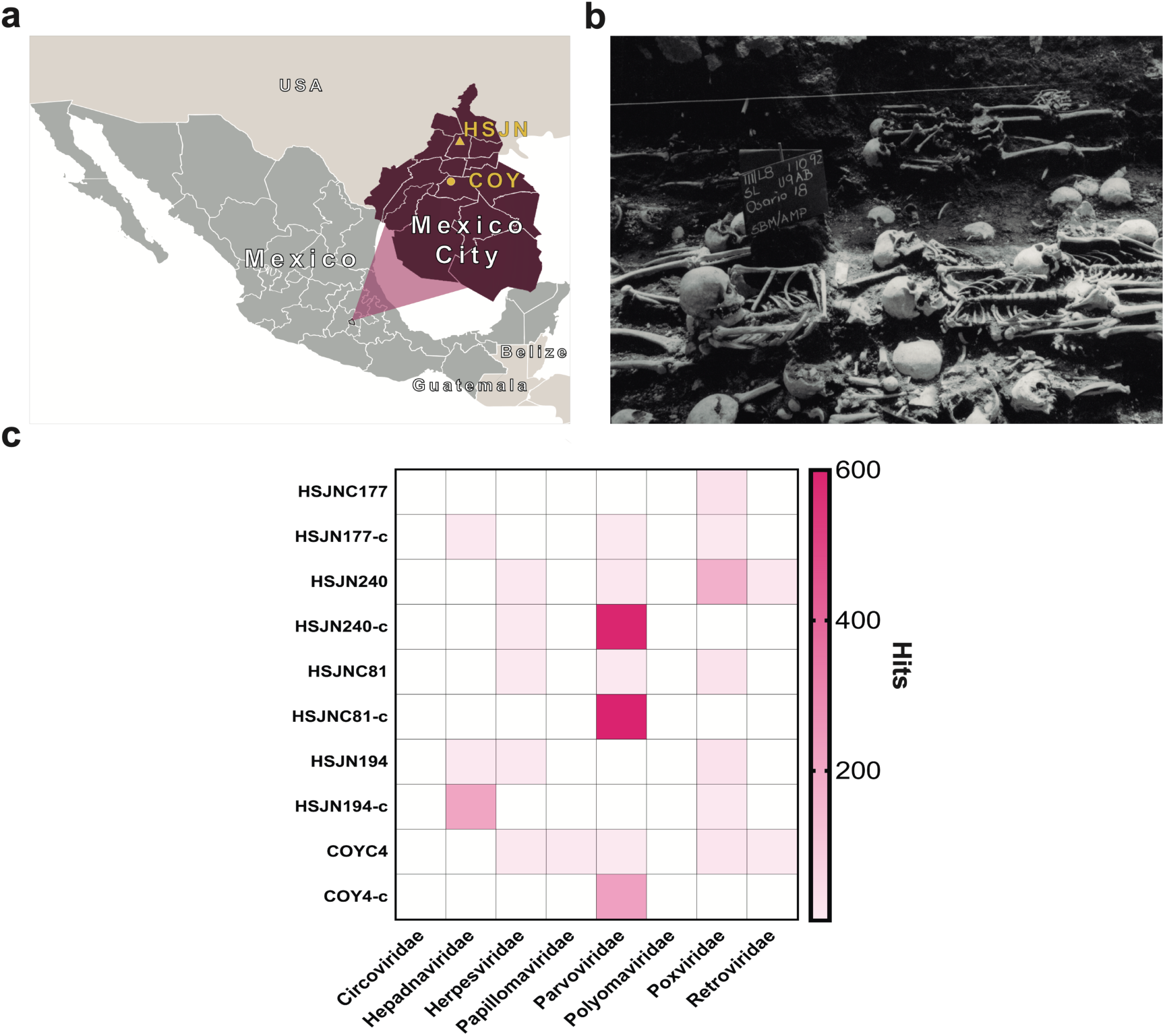
Metagenomic analysis of Colonial individuals reveal HBV-like and B19V-like hits. **a)** Location of the archeological sites used in this study, HSJN (19.431704, -99.141740) is shown as a yellow triangle and COY (19.347079, - 99.159017) as a yellow circle, lines in pink map show current division of Mexico City. **b**) Several individuals discovered in massive burials archaeologically associated with the HSJN and colonial epidemics (photo courtesy of “Secretaria de Cultura INAH, SINAFO, Fototeca DSA”). **c**) Metagenomic analysis performed with MALT 0.4.0 based on the Viral NCBI RefSeq. Viral abundancies were compared and normalized automatically in MEGAN between shotgun (*sample_name*) and capture (*sample_name*-c) NGS data. Only HBV or B19V positive samples are shown (all samples analyzed are shown in Supplementary Figure 2a-b). A capture negative control (HSJN177) is shown.

The study of ancient viral genomes has revealed important insights into the evolution of specific viral families (Barquera et al., 2020; Duggan et al., 2016; Düx et al., 2020; Kahila Bar-Gal et al., 2012; Krause-Kyora et al., 2018, p.; Mühlemann, Jones, et al., 2018, 2018; Mühlemann, Margaryan, et al., 2018; Neukamm et al., 2020; Pajer et al., 2017; Patterson Ross et al., 2018; Xiao et al., 2013), as well as their interaction with human populations (Spyrou et al., 2019). To explore the presence of viral pathogens in circulation during epidemic periods in New Spain, we leveraged the vast historical and archeological information available for the Colonial HSJN. These include the skeletal remains of over 600 individuals recovered from mass burials associated with the hospital’s architectural remnants (Figure 1b). Many of these remains were retrieved from burial contexts suggestive of an urgent and simultaneous disposal of the bodies, as in the case of an epidemic (Meza, 2013; Wesp, 2017). Prior bioarcheological research has shown that the remains of a number of individuals in the HSJN collection displayed dental modifications and/or morphological indicators typical of African ancestry (Meza, 2013), consistent with historical and archeological research that documents the presence of a large number of both free and enslaved Africans and their descendants in Colonial Mexico (Aguirre-Beltrán, 2005). Indeed a recent paleogenomics study reported a Sub-Saharan African origin of three individuals from this collection (Barquera et al., 2020).

Here we describe the recovery and characterization of viral pathogens that circulated in New Spain during Colonial times, using ancient DNA (aDNA) techniques (Supplementary Figure 1). For this work, we sampled skeletal human remains recovered from the HSJN where archeological context suggest victims of epidemics were buried (Meza, 2013) and from “La Concepcion” chapel, one of the first catholic conversion centers in New Spain (Moreno-Cabrera et al., 2015) (Figure 1a). We report the reconstruction of ancient hepatitis B virus (HBV) and human parvovirus B19 (B19V) genomes recovered from these remains. Our findings provide a direct molecular evidence of human viral pathogens of African origin being introduced to New Spain during the transatlantic slave trade.

## RESULTS

We sampled the skeletal remains from two archeological sites, a Colonial Hospital and a Colonial chapel in Mexico City (Figure 1a-b). For the HSJN, 21 dental samples (premolar and molar teeth) were selected based on previous morphometric analyses and dental modifications that suggested an African ancestry (Hernández-Lopez & Negrete, 2012; Karam-Tapia, 2012; Meza, 2013; Ruíz-Albarrán, 2012). The African presence in the Indigenous Hospital might reflect an urgent response to an epidemic outbreak, since hospitals treated patients regardless of the origin of the affected individuals during serious public health crises (Meza, 2013). Dental samples of five additional individuals were selected (based on their conservation state) from “La Concepción” chapel (COY), which is located 10 km south of the HSJN in Coyoacán, a Pre-Hispanic Indigenous neighborhood that became the first Spanish settlement in Mexico City after the fall of Tenochtitlan (Moreno-Cabrera et al., 2015). Following strict aDNA protocols, we processed these dental samples to isolate aDNA for next-generation sequencing (NGS) (Supplementary Figure 1, Methods). Teeth roots (which are vascularized) can be a good source of pathogen DNA(Key et al., 2017), especially in the case of viruses that are widespread in the bloodstream during systemic infection. Accordingly, a number of previous studies have successfully recovered ancient viral DNA from teeth roots (Barquera et al., 2020; Krause-Kyora et al., 2018; Mühlemann, Jones, et al., 2018; Mühlemann, Margaryan, et al., 2018),(Mühlemann et al., 2020).

Metagenomic analysis with MALT (Vågene et al., 2018) (Methods) on the NGS data using the Viral NCBI RefSeq database as a reference (Pruitt et al., 2007), revealed seventeen samples contained at least one normalized hit to viral DNA, particularly similar to *Hepadnaviridae*, *Herpesviridae*, *Parvoviridae* and *Poxviridae* (Figure 1c, Supplementary Figure 2a, Methods). These viral hits revealed the potential to recover ancient viral genomes from these samples. We selected twelve samples for further screening (Figure 1c, Supplementary Figure 2b) based on the DNA concentration of the NGS library and the quality of the hits to a clinically important virus (HBV, B19V, Papillomavirus, Smallpox). To isolate and enrich the viral DNA fraction in the sequencing libraries, biotinylated single-stranded RNA probes designed to capture sequences from diverse human viral pathogens were synthesized (Supplementary Table 1). The selection of the viruses included in the capture design considered the following criteria: 1) DNA viruses previously retrieved from archeological human remains (i.e. Hepatitis B virus, Human Parvovirus B19, Variola Virus), 2) representative viruses from families capable of integrating into the human genome (i.e. *Herpesviridae*, *Papillomaviridae*, *Polyomaviridae*, *Circoviridae*) or 3) RNA viruses with a DNA intermediate (i.e. *Retroviridae*). Additionally, a virus-negative aDNA library, which showed no hits to any viral family included in the capture assay (except for a frequent *Poxviridae*-like region identified as an Alu repeat (Tithi et al., 2018)), was captured and sequenced as a negative control (HSJN177) to estimate the efficiency of our capture assay. Four post-capture libraries had a ∼100-fold increase of HBV-like hits or a ∼50-200-fold increase of B19V-like hits (Figure 1c, Supplementary Table 2) compared to their corresponding pre-capture libraries (Methods). In contrast, the captured negative control (HSJN177) presented a negligible enrichment of these viral hits (Figure 1c, Supplementary Table 2).

We verified the authenticity of the viral sequences by querying the mapped reads against the non-redundant (nr) NCBI database using megaBLAST(Altschul et al., 1990). We only retained reads for which the top hit was to either B19V or HBV, respectively (Supplementary Table 3). 1 To confirm the ancient origin of these viral reads, we evaluated the misincorporation damage patterns using the program mapDamage 2.0 (Jónsson et al., 2013), which revealed an accumulation of C to T mutations towards their 5’ terminal site with an almost symmetrical G to A pattern on the 3’ end (Figure 2a, Supplementary Figure 3a), as expected for aDNA (Briggs et al., 2007). Three ancient B19V genomes were reconstructed (Figure 2b, Supplementary Table 3) with sequence coverages between 92.37% and 99.1%, and average depths of 2.98-15.36X along their single stranded DNA (ssDNA) coding region, which excludes the double stranded DNA (dsDNA) hairpin regions at each end of the genome (Luo & Qiu, 2015). These dsDNA inverse terminal repeats (ITRs) displayed considerably higher depth values (<218X) compared to the coding region consistent with the better *post-mortem* preservation of dsDNA compared to ssDNA (Lindahl, 1993) (Figure 2b). In addition, we reconstructed one ancient HBV genome (Figure 2c, Supplementary Table 3) at 30.8X average depth and with a sequence coverage of 89.9%, including its ssDNA region at a reduced depth (<10X). This genome shows a 6 nucleotide (nt) insertion in the core gene, which is characteristic of the genotype A (Kramvis, 2014). Further phylogenetic analyses (Methods) revealed that the Colonial HBV genome clustered with modern sequences corresponding to sub-genotype A4 (previously named A6) (Pourkarim et al., 2014) (Figure 3a, Supplementary Figure 4). The Genotype A (HBV/GtA) has a broad diversity in Africa reflecting its long history in this continent (Kostaki et al., 2018; Kramvis, 2014), while the sub-genotype A4 has been recovered uniquely from African individuals in Belgium (Pourkarim et al., 2010) and has never been found in the Americas. Regarding the three Colonial B19V genomes from individuals HSJN240, COYC4 and HSJNC81, these were phylogenetically closer to modern B19V sequences belonging to genotype 3 (Figure 3b, Supplementary Figure 5a-b). This B19V genotype is divided into two sub-genotypes: 3a that is mostly found in Africa, and 3b, which is proposed to have spread outside Africa in the last decades (Hübschen et al., 2009). The viral sequences from the individuals HSJN240 and COYC4 are similar to sub-genotype 3b genomes sampled from immigrants (Morocco, Egypt and Turkey) in Germany (Schneider et al., 2008) (Figure 3b, Supplementary Figure 5a-b); while the sequence of the individual HSJNC81 is more similar to a divergent sub-genotype 3a strain (Figure 3b, Supplementary Figure 5a-b) retrieved from a child with severe anemia born in France (Nguyen et al., 1999). These observations support the African origin of the reconstructed colonial viral genomes.

**Figure 2.**
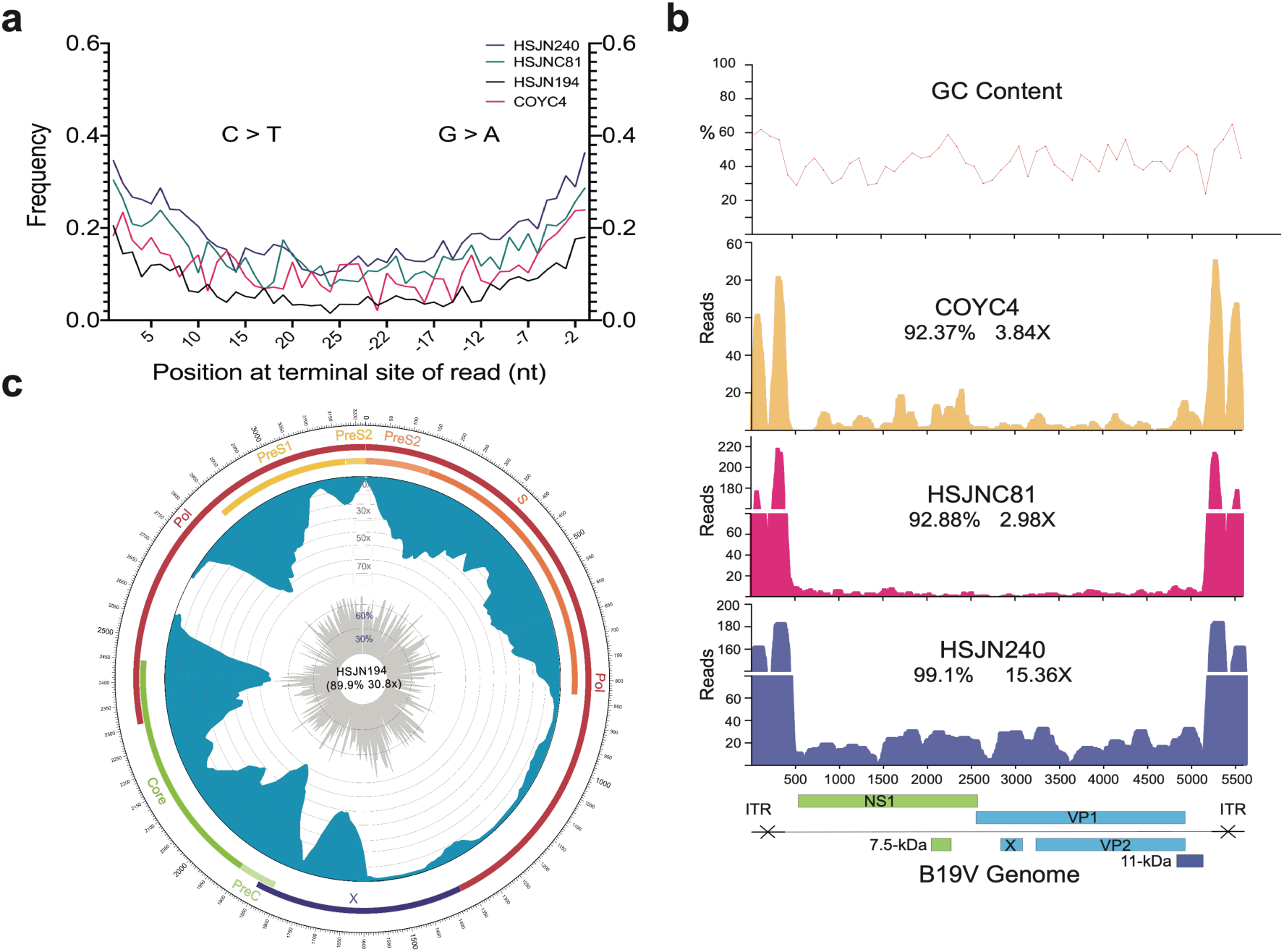
Ancient B19V and HBV ancient genomes. **a**) Superimposed damage patterns of ancient HBV (HSJN194) and B19V (HSJNC81, HSJN240, COYC4), X axis shows the position (nt) on the 5’ (left) and 3’ (right) end of the read, Y axis shows the damage frequency (raw individual damage patterns are shown on Supplementary Figure 3). **b**) B19V ssDNA linear genome, X axis shows position (nt) based on the reference genome (AB550331), and Y axis shows depth (as number of reads), GC content is shown as a percentage of each 100 bp windows, coverage and average depth for the CDS are shown under each individual ID. Schematic of the B19V genome is shown at the bottom. Highly covered regions correspond to dsDNA ITRs shown as crossed arrows. **c**) HBV circular genome, outer numbers show position (nt) based on reference genome (GQ331046), outer bars show genes with names, blue bars represent coverage and gray bars shows GC content each 10 bp windows. Coverage and average depth are shown in the center. Low covered region between S and X overlaps with ssDNA region.

**Figure 3.**
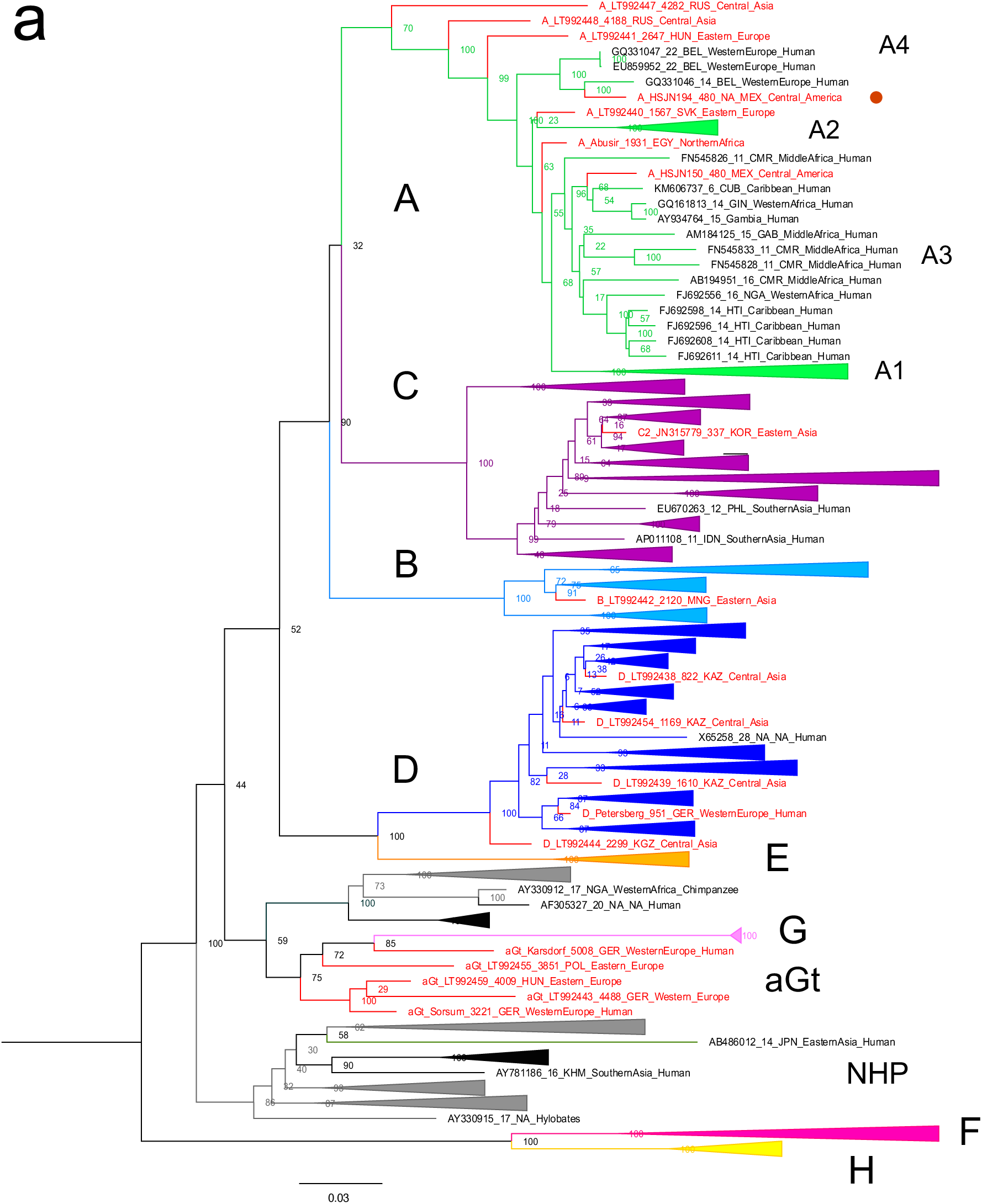
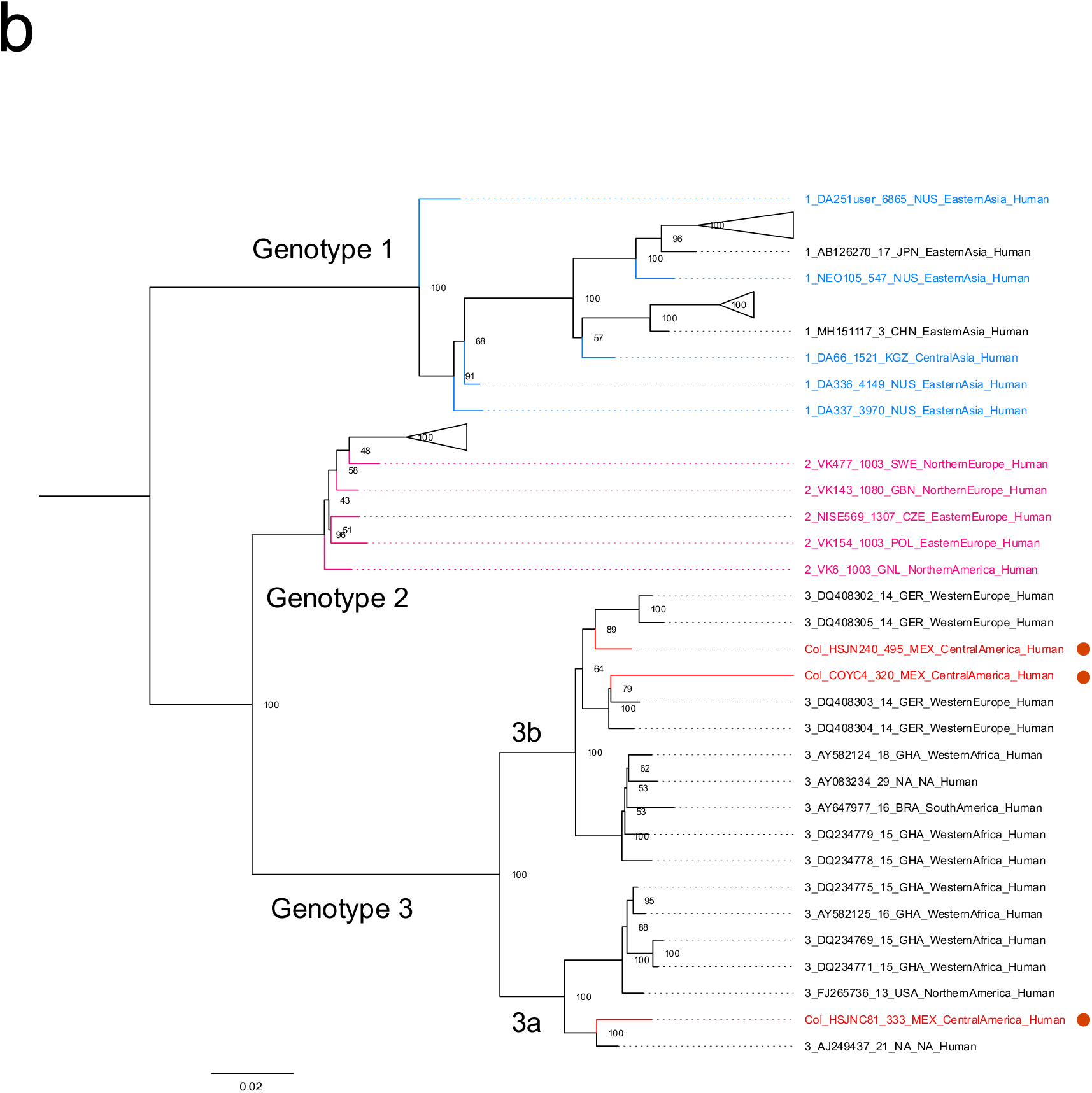
Viral Colonial genomes are similar to modern African genetic diversity. **a-b**. Maximum likelihood trees performed on RAxML 8.2.10 (1000 bootstraps) with a midpoint root, genotypes are named in bold letters and sub-genotypes in italics. Bootstrap values are shown at the node center and triangles represent collapsed sequences from other genotypes. Sequences are named as follows: genotype_ID_sampling.year_country.of.origin_area.of.origin_host. Sequences from this study are highlighted with a red circle on the right. **a)** Based on the HBV whole genome, genotypes are named with letters and each is colored differently, while ancient sequences are shown in red. NHP: non-human primates; **b)** based on B19V CDS where genotypes are named with numbers, and only ancient genomes are colored.

In order to infer the temporal dynamics between our samples and the rest of the viral diversity, we first estimated if the phylogenetic relationships among B19V or HBV genomes had a temporal structure. Similarly to previous studies (Krause-Kyora et al., 2018), we found little or no temporal structure for this HBV phylogeny containing all genotypes (R^2^=0.1351; correlation coefficient=0.3676) (Supplementary Figure 6a-c). The complex evolution of HBV may not be prone to an appropriate genetic dating since multiple recombination and cross-species transmission (Human-Ape) events (Krause-Kyora et al., 2018) occurred throughout its evolution. Since the entire Genotype A has been identified as a recombinant genotype before (Mühlemann, Jones, et al., 2018) we analyzed it independently and identified a stronger temporal signal within this genotype (R^2^=0.722; correlation coefficient=0.8498) (Supplementary Figure 6d-f). In the case of B19V we identified a temporal structure when including all three genotypes (R^2^=0.3837; correlation coefficient=0.6194) (Supplementary Figure 7a-c), in agreement with previous studies (Mühlemann, Margaryan, et al., 2018). Furthermore, we corroborated this temporal structure was not an artifact by a set of tip-dated randomized analyses (Rieux & Balloux, 2016), where none of clock rate 95% highest posterior density (HPD) intervals overlapped with the correctly dated dataset (Supplementary Figure 8).

With these results we then performed a dated coalescent phylogenetic analysis. We inferred a median substitution rate for B19V of 1.03×10^-5^ (95% HPD: 8.66×10^-6^- 1.21×10^-5^) s/s/y under a strict clock and a constant population prior and a substitution rate of 2.62×10^-5^ (95% HPD: 1.50×10^-5^-3.98×10^-5^) s/s/y under a relaxed log normal clock and a constant population prior. The divergence times from the most recent common ancestor of genotypes 1, 2 and 3 under a strict clock were 7.19 (95% HPD: 6.98-7.46), 2.11 (95% HPD: 1.83-2.51), and 3.64 (95% HPD: 3.04-4.33) ka, respectively. The inferred substitution rates and divergence times from the most recent common ancestor for genotypes 1 and 2 were similar to previous estimations (Mühlemann, Margaryan, et al., 2018) that included much older sequences, while the divergence of genotype 3 was subtly older since no other ancient genotype 3 had been reported previously.

Next, we used the *de novo* generated sequence data to determine the mitochondrial haplogroup of the hosts, as well as their autosomal genetic ancestry using the 1000 Genomes Project (1000 Genomes Project Consortium et al., 2015) as a reference panel (Figure 4a, Supplementary Table 4). The nuclear genetic ancestry analysis showed that all three HSJN individuals, from which the reconstructed viral genomes were isolated, fall within African genetic variation in a Principal Component Analysis plot (Figure 4a), while their mitochondrial aDNA belong to the L haplogroup, which has high frequency in African populations (Supplementary Table 4, Supplementary Figure 3b). Additionally, we performed ^87^Sr/^86^Sr isotopic analysis on two of the HSJN individuals using teeth enamel as well as phalange (HSJN240) or parietal bone (HSJNC81) to provide insights on the places of birth (adult enamel) and where the last years of life were spent (phalange/parietal). The ^87^Sr/^86^Sr ratios measured on the enamel of the individual HSJNC81 (0.71098) and HSJN240 (0.71109) are similar to average ^87^Sr/^86^Sr ratios found in soils and rocks from West Africa (average of 0.71044, Supplementary Figure 9, Supplementary Tables 6 and 7), as well as to ^87^Sr/^86^Sr ratios described in first generation Africans in the Americas (Barquera et al., 2020; Bastos et al., 2016; Fricke et al., 2020; T. D. Price et al., 2012; Schroeder et al., 2009). In contrast, the ^87^Sr/^86^Sr ratios on the parietal and phalange bones from the HSJNC81 (0.70672) and HSJN240 (0.70755), show lower values similar to those observed in the Trans Mexican Volcanic Belt where the Mexico City Valley is located (0.70420 - 0.70550, Supplementary Figure 9, Supplementary Tables 6 and 7). Moreover, radiocarbon dating of HSJN240 (1442-1608 CE, years calibrated for 1σ) and HSJN194 (1472-1625 CE, years calibrated for 1σ) (Supplementary Table 4, Supplementary Figure 10) indicates that these individuals arrived during the first decades of the Colonial period, when the number of enslaved individuals arriving from Africa was particularly high (Aguirre-Beltrán, 2005). Strikingly, Colonial individual COYC4, who was also infected with an African B19V strain, clusters with present-day Mexicans and Peruvians from the 1000 Genomes Project (Figure 4a). An ADMIXTURE (Alexander & Lange, 2011) analysis with these data confirmed a predominant Native American and African genetic component (Figure 4b), as expected for a post-contact individual. The B19V ancient genome from the individual COYC4 is the first genotype 3 genome obtained from a non-African individual and suggests that following the introduction from Africa, the virus (B19V) spread and infected people of different ancestries during Colonial times.

**Figure 4.**
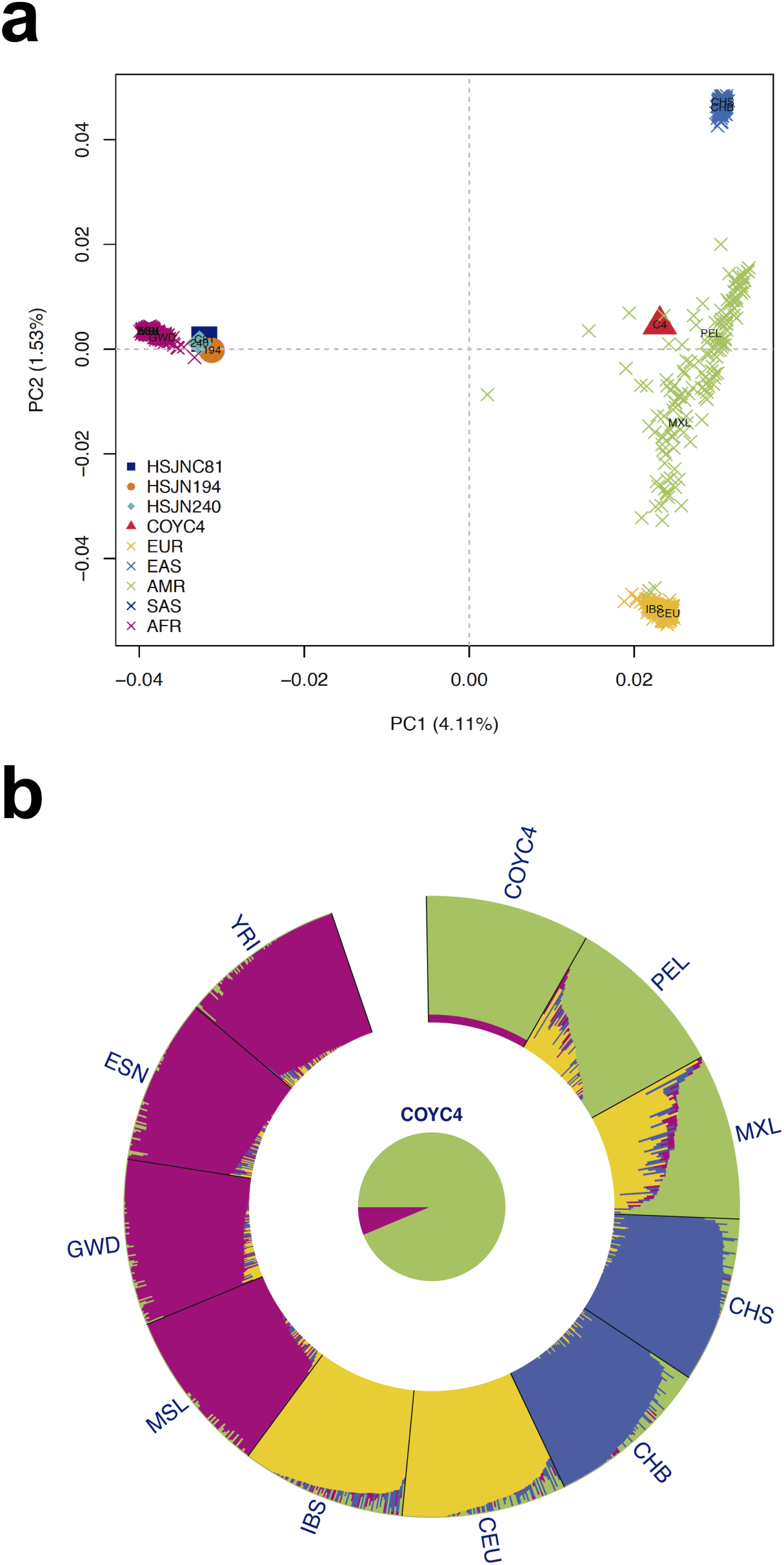
Human hosts are similar to modern African genetic diversity. **a)** PCA showing genetic affinities of ancient human hosts compared to the 1000 Genomes Project reference panel. Crosses (X) show individuals from the reference panel while other shapes show human hosts from which ancient HBV (HSJN194) and B19V (HSJNC81, HSJN240, COYC4) sequences were recovered. Clusters are colored in five super populations EUR: Europeans (IBS, CEU); EAS: East Asian (CHB); AMR: Ad Mixed Americans (MXL, PEL); SAS: South Asians (CHS) and AFR: Africans (YRI, ESN, GWD, MSL). Three letter code is based on the 1000 Genomes Project nomenclature. **b)** Admixture analysis with COYC4 intersected sites with 1000 Genomes MEGA array, run with k=4 for 100 replicates. Each color shows a different component using the same colors as in the PCA. In the center a pie chart shows the proportion of native American (green) and African (magenta) genetic components.

## DISCUSSION

In this study we reconstructed one HBV and three B19V ancient genomes from four different individuals using NGS, metagenomics and in-solution targeted enrichment methods (Figure 2b, c, Supplementary Figure 1). Several lines of evidence support the ancient nature of these viral sequences, in contrast to environmental contamination or a capture artifact. First, our negative control was not enriched for B19V or HBV hits in our capture sequencing (Figure 1c). For those samples that showed an enrichment in viral sequences after capture, the reads covered the reference genomes almost in their entirety and displayed deamination patterns at the terminal ends of the reads, as expected for aDNA (Figure 2a). Moreover, it is important to notice that B19V and HBV are blood-borne human pathogens that are not present in soil or the environment, and that DNA from these viruses had never been extracted before in the aDNA facilities used for this study.

We also described an unusual coverage pattern on the B19V genome, where the dsDNA hairpins at its terminal sites are highly covered reflecting a better stability of these regions over time (Figure 2b). Similarly, the partially circular dsDNA genome from HBV was poorly covered at the ssDNA region (Figure 2c), as found in three previously reported ancient HBV genomes (Krause-Kyora et al., 2018) (Supplementary Discussion 1). The variable read coverage in both viruses argues against an integration event of these viruses, which would result in a uniform dsDNA coverage; further analyses are needed to determine if the aDNA retrieved in this and other studies comes from systemic circulating virions or from systemic cell-free DNA intermediates (Cheng et al., 2019) produced after viral replication in the bone marrow or liver, for B19V and HBV, respectively (Broliden et al., 2006; Yuen et al., 2018).

The ancient B19V genomes were assigned to genotype 3. This genotype is the most prevalent in West Africa (Ghana: 100%, n=11; Burkina Faso: 100%, n=5) (Candotti et al., 2004; Hübschen et al., 2009; Rinckel et al., 2009) and a potential African origin has been suggested (Candotti et al., 2004). It has also been sporadically found outside Africa (Jain et al., 2015),(Candotti et al., 2004; Rinckel et al., 2009) in countries historically tied to this continent, like Brazil (50%, n=12) (Freitas et al., 2008; Sanabani et al., 2006), India (15.4%, n=13) (Jain et al., 2015), France (11.4%, n=79) (Nguyen et al., 1999; Servant et al., 2002), and USA (0.85%, n=117) (Rinckel et al., 2009) as well as in immigrants from Morocco, Egypt, and Turkey in Germany (6.7%, n=59) (Schneider et al., 2008). Two other genotypes, 1 and 2 exist for this virus. Genotype 1 is the most common and is found worldwide, while the almost extinct genotype 2 is mainly found in elderly people from Northern Europe (Pyöriä et al., 2017). Ancient genomes from genotypes 1 and 2 have been recovered from Eurasian samples, including a genotype 2 B19V genome from a 10^th^ century Viking burial in Greenland (Mühlemann, Margaryan, et al., 2018). ^87^Sr/^86^Sr isotopes on individuals from such burial revealed they were immigrants from Iceland (Mühlemann, Margaryan, et al., 2018), suggesting an introduction of the genotype 2 to North America during Viking explorations of Greenland.

While serological evidence indicates that B19V currently circulates in Mexico, only the presence of genotype 1 has been formally described using molecular analyses (Valencia Pacheco et al., 2017). Taken together, our results are consistent with an introduction of the genotype 3 to New Spain as a consequence of the transatlantic slave trade imposed by the European colonization. This hypothesis is supported by the ^87^Sr/^86^Sr isotopic analysis, which gives evidence that the individuals from the HSJN with B19V (HSJN240, HSJNC81) were born in West Africa and spent their last years of life in New Spain (Supplementary Figure 9). Furthermore, the radiocarbon ages of individuals HSJN240, HSJN194 (Supplementary Figure 10) support this notion as they correspond to the Early Colonial period, during which the number of enslaved Africans arriving was higher compared to later periods (Aguirre-Beltrán, 2005). Remarkably, a B19V genome belonging to the genotype 3 was recovered from an admixed individual (COYC4) (Figure 4b) with a predominant Indigenous ancestry as well as some African. COY4 was excavated in an independent archeological site 10 Km south of the HSJN (Figure 1a), supporting the notion that viral transmissions between African individuals and people of different ancestries occurred during the Colonial period in Mexico City.

The genotype A from HBV is highly diverse in Africa, reflecting its long evolutionary history, and likely originated somewhere between Africa, Middle East and Central Asia (Kostaki et al., 2018). The introduction of the genotype A from Africa to the Americas has been proposed based on phylogenetic analysis of modern strains from Brazil (Freitas et al., 2008; Kostaki et al., 2018) and Mexico (Roman et al., 2010), and more precisely to the sub-genotype A1 using sequences from Martinique (Brichler et al., 2013), Venezuela (Quintero et al., 2002), Haiti (Andernach et al., 2009), and Colombia (Alvarado-Mora et al., 2012). Recently, a similar introduction pattern was proposed for the quasi genotype A3 based on an ancient HBV genome recovered from an ancient African individual sampled in Mexico (Barquera et al., 2020). The origin of the sub-genotype 4 is controversial with an apparent African origin based on sequences recovered from African individuals in Europe (Pourkarim et al., 2010). The Colonial ancient HBV genome reconstructed in our work is assigned to genotype A4 (Figure 3a, Supplementary Figure 4), which represents the first report of this sub-genotype in the Americas and further supports its African origin. The introduction of the pathogens from Africa to the Americas has been proposed for other human-infecting viruses such as smallpox (Mandujano-Sánchez et al., 1982; Somolinos d’Árdois, 1982), based on historical records; or Yellow fever virus (Bryant et al., 2007), HTLMV-1 (Gadelha et al., 2014), Hepatitis C virus (genotype 2) (Markov et al., 2009) and human herpes simplex virus (Forni et al., 2020) based on phylogenetic analysis of modern strains from Afro-descendant or admixed human populations.

Although we cannot assert where exactly the African-born individuals in this study contracted B19V or HBV (Africa, America, or the Middle Passage) nor if the cause of their deaths can be attributed to such infections, the identification of ancient B19V and HBV in contexts associated with Colonial epidemics in Mexico City is still relevant in light of their paleopathological marks and the clinical information available for the closest sequences in the phylogenetic analyses. The reconstructed ancient B19V genome from individual HSJNC81 is closest to the V9 strain, which was isolated from an individual with severe anemia(Nguyen et al., 1999) (AJ249437) (Figure 3b). Noteworthy, individual HSJNC81 displayed cribra orbitalia in the eye sockets and porotic hyperostosis on the cranial vault (Supplementary Figure 11), morphological changes typically associated with anemias of varying different causes (Angel, 1966). It is acknowledged that B19V infection can cause severe or even fatal anemia due to the low level of hemoglobin when present simultaneously with other blood disorders, as thalassemia, sickle-cell anemia, malaria and iron deficiency (Broliden et al., 2006; Heegaard & Brown, 2002). Therefore, since B19V infects precursors of the erythroid lineage (Broliden et al., 2006), it is possible that the morphological changes found in HSJNC81 might be the result of a severe anemia caused or enhanced by a B19V infection (Supplementary Discussion 2). Moreover, the identification of ancient B19V in a Colonial context is noteworthy considering several recent reports that reveal that measles-like cases were actually attributable to B19V (De Los Ángeles Ribas et al., 2019; Rezaei et al., 2016). Therefore it is possible that B19V might have been responsible for some of the numerous cases of measles that were described in early 16^th^ century Mexico (Acuña-Soto et al., 2004; Mandujano-Sánchez et al., 1982; Wesp, 2017), as well as in historical records that account for the treatment of an outbreak of measles at the HSJN in 1531 (Meza, 2013) (Supplementary Discussion 2). Nevertheless, this hypothesis requires additional comprehensive studies aimed to characterize the presence of measles and rubella viruses from ancient remains, a task that imposes difficult technical challenges given that RNA is known to degrade rapidly. In fact most ancient viral RNA genomes have been recovered only from formalin-fixed tissue (Düx et al., 2020; Xiao et al., 2013)

Furthermore, historical records of the autopsies of victims of the 1576 *Cocoliztli* epidemic treated at the HSJN, describe the observation of enlarged hard liver and jaundice (Acuña-Soto et al., 2002, 2004; Malvido & Viesca, 1982; Marr & Kiracofe, 2000; Somolinos d’Árdois, 1982), which could be explained by severe liver damage or epidemic hepatitis (Acuña-Soto et al., 2004; Malvido & Viesca, 1982). This is noteworthy given both viruses HBV and B19V proliferate in the liver and are associated to hepatitis and jaundice (Broliden et al., 2006; Yuen et al., 2018). However, it is important to acknowledge that both viruses have also been previously identified in aDNA datasets not necessarily associated with disease or epidemic contexts (Kahila Bar-Gal et al., 2012; Krause-Kyora et al., 2018; Mühlemann, Jones, et al., 2018; Patterson Ross et al., 2018), thus establishing a direct link would require additional samples and a more comprehensive pathogen screening to rule out the involvement of other pathogens. Finally, although our data does not provide conclusive evidence of the involvement of HBV and B19V in the reported manifestations of liver damage in *Cocoliztli* autopsies, the identification of these viruses in likely victims of epidemic outbreaks in the Colonial period opens up new opportunities for investigating the presence of these viruses in similar contexts. This type of research is particularly relevant when considering previous hypotheses favoring the synergistic action of different types of pathogens in these devastating Colonial epidemics (Somolinos d’Árdois, 1982) (Supplementary Discussion 3).

It is important to emphasize that our findings should be interpreted with careful consideration of the historical and social context of the transatlantic slave trade. This cruel episode in history involved the forced displacement of millions of individuals to the Americas (ca. 250,000 to New Spain (Aguirre-Beltrán, 2005)) under inhumane, unsanitary and overcrowded conditions that, with no doubt, favored the spread of infectious diseases (Mandujano-Sánchez et al., 1982). Therefore, the introduction of these and other pathogens from Africa to the Americas should be attributed to the brutal and harsh conditions of the Middle passage that enslaved Africans were subjected to by traders and colonizers, and not to the African peoples themselves. Moreover, the adverse life conditions for enslaved Africans and Native Americans, especially during the first decades after colonization, surely favored the spread of diseases and emergence of epidemics (Mandujano-Sánchez et al., 1982). Integrative and multidisciplinary approaches are thus needed to understand this phenomenon at its full spectrum.

In summary, our study provides direct aDNA evidence of HBV and B19V introduced to the Americas from Africa during the transatlantic slave trade. The isolation and characterization of these ancient HBV and B19V genomes represent an important contribution to the only ancient viral genome recently reported in the Americas (Barquera et al., 2020). Our results expand our knowledge on the viral agents that were in circulation during Colonial epidemics like *Cocoliztli*, some of which resulted in the catastrophic collapse of the immunologically-naïve Indigenous population. Although we cannot assign a direct causality link between HBV and B19V and *Cocoliztli*, our findings confirm that these potentially harmful viruses were indeed circulating in individuals found in archeological contexts associated with this epidemic outbreak. Further analyses from different sites and samples will help understand the possible role of these and other pathogens in Colonial epidemics, as well as the full spectrum of pathogens that were introduced to the Americas during European colonization.

## METHODS

### Sample selection and DNA extraction

Dental samples (premolars and molars) were obtained from twenty-one individuals from the skeletal collection of the HSJN, selected based on their African-related skeletal indicators (Hernández-Lopez & Negrete, 2012; Karam-Tapia, 2012; Meza, 2013; Ruíz-Albarrán, 2012). Five additional samples were taken from “La Concepción” chapel, based on their conservation state. Permits 401.1S.3-2018/1373 and 401.1S.3-2020/1310 to carry out this sampling and aDNA analyses were obtained by the Archeology Council of the National Institute of Anthropology and History (INAH) for the Hospital San Jose de los Naturales and “La Concepción” chapel, respectively.

### DNA extraction and NGS library construction

Bone samples were transported to a dedicated ancient DNA clean-room laboratory at the International Laboratory for Human Genome Research (LIIGH-UNAM, Querétaro, Mexico), were DNA extraction and NGS-libraries construction was performed under the guidelines on contamination control for aDNA studies (Warinner et al., 2017). Previously reported aDNA extraction protocols were used for the HSJN (Dabney et al., 2013) and COY (Rohland & Hofreiter, 2007) samples. Double-stranded DNA (dsDNA) indexed (6bp) sequencing libraries were constructed from the DNA extract, as previously reported (Meyer & Kircher, 2010). In order to detect contaminants in reagents or by human manipulation, extraction and library constructions protocols included negative controls (NGS blanks) that were analyzed in parallel with the same methodology. The resulting NGS dsDNA indexed libraries were quantified with a Bioanalyzer 2100 (Agilent) and pooled into equimolar concentrations.

### NGS sequencing

Pooled libraries were paired-end sequenced on an Illumina NextSeq550 at the “Laboratorio Nacional de Genómica para la Biodiversidad” (LANGEBIO, Irapuato, Mexico), with a Mid-output 2×75 format. The reads obtained (R1 and R2) were merged (>11bp overlap) and trimmed with AdapterRemoval 1.5.4 (Schubert et al., 2016). Overlapping reads (>30 bp in length) were kept and mapped to the human genome (hg19) using BWA 0.7.13 (Heng Li & Durbin, 2009). Mapped reads were used for further human analysis (genetic ancestry, and mitochondrial haplogroup determination), whereas unmapped reads were used for metagenomic analysis and viral genome reconstruction.

### Metagenomic analyses

The NCBI Viral RefSeq database was downloaded on February 2018; this included 7530 viral genomes. MALT 0.4.0 (Vågene et al., 2018) software was used to taxonomically classify the reads using the viral genomes database. The viral database was formatted automatically with malt-build once, and not human (unmapped) reads were aligned with malt-run (85 minimal percent identity). The produced RMA files with viral abundances were normalized based on the smallest sample size (default) and compared to all the samples from the same archeological site with MEGAN 6.8.0 (Huson et al., 2016).

### Capture-enrichment Assay

Twenty-nine viruses were included in the in-solution enrichment design, the complete list of NCBI IDs is provided in Supplementary methods 5 and Supplementary Table 1. It contained viral genomes previously recovered from archeological remains like B19V, B19V-V9, and HBV (consensus genomes), selected VARV genes, as well as clinically important viral families that are able to integrate into the human genome, have dsDNA genomes, or dsDNA intermediates. The resulting design comprised 19,147 ssRNA 80 nt probes targeting, with a 20 nt interspaced distance, the whole or partial informative regions of eight viral families of clinical relevance (*Poxviridae, Hepadnaviridae, Parvoviridae, Herpesviridae, Retroviridae, Papillomaviridae, Polyomaviridae, Circoviridae*). To avoid a simultaneous false-positive DNA enrichment, low complexity regions and human-like (hg38) sequences were removed (*in sillico*). The customized kit was produced by Arbor Biosciences (Ann Arbor, MI, USA). Capture-enrichment was performed on the indexed libraries based on the manufacturer’s protocol (version 4) to pull-down aDNA with minimal modifications. Libraries were amplified with 18-20 cycles (Phusion U Hot Start DNA Polymerase by Thermo Fischer Scientific), purified with SPRISelect Magnetic Beads (Beckman Coulter) and quantified with a Bioanalyzer 2100 (Agilent). Amplified libraries were then pooled in different concentrations and deep sequenced yielding >1×10^6^ non-human reads (Supplementary Table 4) in order to saturate the target viral genome. Reads generated from each enriched library were analyzed exactly as the shotgun (not-enriched) libraries. Normalized abundances between shotgun and captured libraries were compared in MEGAN 6.8.0 (Huson et al., 2016) to evaluate the efficiency and specificity of the enrichment assay.

### Viral datasets

The full list of accession numbers of the following datasets is given in Supplementary Methods 8.

HBV-Dataset-1 (HBV/DS1): comprises 38 HBV genomes from modern A-J human genotypes, 2 well-covered ancient HBV genomes (LT992443, LT992459) and a wholly monkey genome.

HBV-Dataset-2 (HBV/DS2): comprises 593 whole genomes downloaded from the NCBI database in August 2020, that included genomes from A-J genotypes as well as non-human primates HBV genomes (gibbon, gorilla, and chimpanzee), 19 ancient HBV genomes (Barquera et al., 2020; Kahila Bar-Gal et al., 2012; Krause-Kyora et al., 2018, p.; Mühlemann, Jones, et al., 2018; Neukamm et al., 2020; Patterson Ross et al., 2018) and one ancient HBV genome from this study (HSJN194).

B19V-Dataset-1 (B19V/DS1): comprises 13 B19V genomes from human genotypes 1-3 as well as a bovine parvovirus.

B19V-Dataset-2 (B19V/DS2): comprises 109 genomes from 1 to 3 B19V genotypes downloaded from the NCBI database in August 2020, that included the 10 best-covered ancient genomes from genotype 1 and 2 (Mühlemann, Margaryan, et al., 2018) as well as 3 ancient B19V from this study. Since many of the reported genomes in our dataset are not complete, only the whole coding region (CDS) was used.

### Genome Reconstruction and authenticity

#### HBV

Non-human reads were simultaneously mapped to HBV/DS1 with BWA (aln algorithm) with seedling disabled (Schubert et al., 2012). The reference sequence with the most hits was used to map uniquely to this reference and generate a BAM alignment without duplicates (Ref: GQ331046), from which damage patterns were determined and damaged sites rescaled using mapDamage 2.0 (Jónsson et al., 2013), the rescaled alignment was used to produce a consensus genome. All the HBV mapped reads were analyzed through megaBLAST (Altschul et al., 1990) using the whole NCBI nr database, in order to verify they were assigned uniquely to HBV (carried out with Krona 2.7 (Ondov et al., 2011)).

#### B19V

The reconstruction of the B19V ancient genome was done as previously reported from archeological skeletal remains (Mühlemann, Margaryan, et al., 2018), but increasing the stringency of some parameters as described next. Non-human reads were mapped against B19V/DS1 with BWA (aln algorithm) with seedling disabled(Schubert et al., 2012), if more than 50% of the genome was covered, the sample was considered positive to B19V. Reads from the B19V-positive libraries were aligned with blastn (-evalue 0.001) to B19V/DS1 in order to recover all the parvovirus-like reads. To avoid local alignments, only hits covering >85% of the read were kept and joined to the B19V mapped reads (from BWA), duplicates were removed. The resulting reads were analyzed with megaBLAST (Altschul et al., 1990) using the whole NCBI nr database to verify the top hit was to B19V (carried out with Krona 2.7 (Ondov et al., 2011)). This pipeline was applied for two independent enrichments assays per sample and the filtered reads from the two capture rounds were joined. The merged datasets per sample were mapped using as a reference file the three known B19V genotypes with GeneiousPrime 2019.0.4 (Kearse et al., 2012) using median/fast sensibility and iterate up to 5 times. The genotype with the longest covered sequence was selected as the reference for further analysis (Ref: AB550331). Deamination patterns for HBV and B19V were determined with mapDamage 2.0 (Jónsson et al., 2013) and damaged sites were rescaled in the same program to produce a consensus whole genome using SAMtools 1.9 (H. Li et al., 2009).

### Phylogenetic analyses

HBV/DS2 and B19V/DS2 were aligned independently in Aliview (Larsson, 2014) (Muscle algorithm), and evolutionary models were tested under an AICc and BIC in jModelTest (Darriba et al., 2012). A neighbor joining tree with 1000 bootstraps was generated in MEGA (Kumar et al., 2018) using a number of differences model for both alignments. A maximum likelihood tree with 1000 bootstraps was produced in RAxML 8.2.10 (Stamatakis, 2014) using a Generalized Time-Reversible (GTR)+G model for both B19V and HBV.

To test the presence of a temporal structure in our DS2 (HBV and B19V) a root-tip distance analysis was performed on Tempest 1.5.3 (Rambaut et al., 2016), a more detailed analysis was also carried out on a subset (HBV/DS2.1) of sequences just containing genomes from the HBV Genotype A.

The temporal structure found in the B19V/DS2 with the root-tip distance analysis was corroborated by a date randomization test (DRT) using 20 combinations of our B19V/DS2 with TipDatingBeast 1.0.5 (Rieux & Khatchikian, 2017) and BEAST 2.5.1 (Drummond et al., 2012).

Since our B19V/DS2 presented a clock-like structure with both methods, a dated Bayesian analysis was generated to estimate the impact of the Colonial viral genomes on the divergence time from the most recent common ancestor (MRCA). We used BEAST 2.5.1 (Drummond et al., 2012), with a relaxed lognormal or strict molecular clock with constant population, Bayesian skyline population or coalescent exponential population prior. All parameters were mixed and converged into an estimated sample size (ESS) >200 analyzed in Tracer 1.7 (Rambaut et al., 2018). The first 25% of trees where discarded (burn in) and a Maximum Clade Credibility Tree was created with TreeAnnotator (Drummond et al., 2012). The generated trees were visualized and edited in FigTree 1.4.3 with a midpoint root.

### Human population genetic analyses

Human-mapped reads (BWA aln) obtained from the pre-capture sequence data of viral-positive samples were used to infer the genetic ancestry of the hosts. A Principal Components Analysis (PCA) was carried out using 10 populations (IBS: Iberian from Spain; CEU: Utah Residents with Northern and Western European Ancestry; CHB: Han Chinese in Beijing; MXL: Mexican Ancestry from Los Angeles; PEL: Peruvians from Lima; CHS: Southern Han Chinese; YRI: Yoruba in Ibadan; ESN: Esan in Nigeria; GWD: Gambian; MSL: Mende in Sierra Leone) from the 1000 Genomes Project (1000 Genomes Project Consortium et al., 2015) reference panel including genotype data of 1,562,771 single nucleotide variants (SNVs) present in the MEGA array (Wojcik et al., 2019) from 2,504 individuals (phase 3). Genomic alignments of each ancient individual (HSJNC81, HSJN240, HSJN194 and COYC4) were intersected with the positions of the SNVs present in the reference panel genotype data. Pseudo haploid genotypes were called by randomly selecting one allele at each intersected site and filtering by a base quality >30. Pseudo haploid genotypes were also called for the complete reference panel. PCA was performed on the merged ancient and modern dataset with smartpca (EIGENSOFT package) (Patterson et al., 2006; A. L. Price et al., 2006) using the option *lsqproject* to project the ancient individuals into the PC space defined by the modern individuals.

### Ancestry composition of individual COYC4

A total of 1,246 sites intersected between the 1000 Genomes Project reference panel and COYC4 ancient genome (see previous section for details). The program ADMIXTURE (Alexander & Lange, 2011) was run with K values of 2 to 5 and 100 replicates for each K using a different seed number. For each K, the ADMIXTURE run with the best likelihood was chosen to plot it using AncestryPainter (Feng et al., 2018).

### Mitochondrial haplogroup and sex determination

NGS reads were mapped to the human mitochondrial genome reference (rCRS) with BWA (aln algorithm, -l default), the alignment file was then used to generate a consensus mitochondrial genome with the program Schmutzi(Renaud et al., 2015) The assignment of the mitochondrial haplogroup was carried out with Haplogrep (Kloss-Brandstätter et al., 2011; Weissensteiner et al., 2016) using the consensus sequence as the input. Assignment of biological sex was inferred based on the fraction of reads mapped to the Y-chromosome (Ry) compared to those mapping to the Y and X-chromosome (Skoglund et al., 2013). Ry<0.016 and Ry>0.075 were considered XX or XY genotype, respectively. The resulting sex was coherent with the one inferred morphologically (Supplementary Methods).

## ACKNOWLEDGMENTS

This work was funded by the Wellcome Trust Sanger grant number 208934/Z/17/Z, and by project IA201219 PAPIIT-DGAPA-UNAM. D.B-M is an Open Philanthropy Fellow of the Life Sciences Research Foundation (LSRF). We thank the INAH Archeology Council for the sample permissions 401.1S.3-2018/1373 and 401.1S.3-2020/1310 for this study using samples from the Hospital San Jose de los Naturales and the Temple of Immaculate Conception (La Conchita), respectively. We are grateful with Teodoro Hernández Treviño, Gerardo Arrieta García from the “Laboratorio Universitario de Geoquímica Isotópica” (LUGIS-UNAM) for their technical support in performing the ^87^Sr/^86^Sr analyses and to Luis Alberto Aguilar Bautista, Alejandro de León Cuevas, Carlos Sair Flores Bautista and Jair Garcia Sotelo from the “Laboratorio Nacional de Visualización Científica Avanzada” (LAVIS/UNAM) who stored our data and provided the computational resources to perform this study. We thank Alejandra Castillo Carbajal and Carina Uribe Díaz for technical support throughout the project.

## DATA AVAILABILITY

Reconstructed genomes from this study are available in Genbank under accession number MT108214, MT108215, MT108216, MT108217. Accession number of sequences used in phylogenetic analysis are indicated in supplementary information. NGS data is available upon reasonable request.

## Notes

### Competing Interest Statement

The authors have declared no competing interest.

