## Supplementary_file for "Ancient viral genomes reveal introduction of HBV and B19V into Mexico during the transatlantic slave trade"

### Supplementary Information

Axel A. Guzmán-Solís[1,2]*, Daniel Blanco-Melo[3]*‡, Viridiana Villa-Islas[1], Miriam J. Bravo-López[1], Marcela Sandoval-Velasco[4], Julie K. Wesp[5], Jorge A. Gómez-Valdés[6], María de la Luz Moreno-Cabrera[7], Alejandro Meraz-Moreno[7], Gabriela Solís-Pichardo[8], Peter Schaaf[9], Benjamin R. tenOever[3] & María C. Ávila-Arcos[1]‡.

1.International Laboratory for Human Genome Research, Universidad Nacional Autónoma de México (México); 2.Aaron Diamond AIDS Research Center, Columbia University Vagelos College of Physicians and Surgeons (NY, USA); 3.Department of Microbiology, Icahn School of Medicine at Mount Sinai (NY, USA); 4.Section for Evolutionary Genomics, The Globe Institute, Faculty of Health, University of Copenhagen (Denmark); 5.Department of Sociology and Anthropology, North Carolina State University (USA); 6.Escuela Nacional de Antropología e Historia (México); 7.Instituto Nacional de Antropología e Historia (México); 8.Laboratorio Universitario de Geoquímica Isotópica (LUGIS), Instituto de Geología, Universidad Nacional Autónoma de México (México); 9.LUGIS, Instituto de Geofísica, Universidad Nacional Autónoma de México (México).

CONTENTS:


- **Supplementary Methods 1-12**
- **Supplementary Discussion 1-3**
- **Supplementary Tables 1-7**
- **Supplementary Figures 1-11**
- **Supplementary References**

**Supplementary methods:**

1. Archeological context of the HSJN samples

The individuals analyzed in this work are part of a skeletal collection associated with the Hospital Real San José de los Naturales (HSJN), a Colonial hospital established in 1531 in what is presently the historic downtown of Mexico City (Muriel, 1956) (Figure 1a). The HSJN was established in the Americas after the arrival of Europeans (J. K. Wesp, 2017), and it was the first royally-sponsored hospital specifically designated to care for the local Indigenous population (Rodríguez-Sala, 2005; Zedillo, 1984).

During the construction of a new line for the Mexico City Metro system in the early 1990s, a team of archeologists uncovered the architectural remains of the HSJN (Báez-Molgado & Meza, 1995), as well as a large collection of skeletal remains that included both articulated skeletons and isolated crania corresponding to a total of over 600 individuals (Báez-Molgado & Meza, 1995) (Figure 1b).

Two of the individuals from whom the ancient viral genomes were retrieved (HSJN194 and HSJN240) are mostly complete articulated skeletons and one individual (HSJNC81) is an isolated cranium recovered during the early excavation stage and does not have any associated postcranial elements. The archeologists suggested that they were deposited during an infectious disease epidemic (Figure 1b) (Cabrera-Torres & García-Martínez, 1998).

HSJN194 (Supplementary Figure 11) is a middle adult, morphologically male (coherent with inferred genetic sex) (Supplementary Table 4), between 35–49 years old with moderate amounts of biomechanical stress, including osteoarthritic bone growth on the upper and lower limbs. He has moderate amounts of dental wear and dental calculus (calcified plaque), as well as a small amount of periodontal disease with at least two maxillary teeth that were lost *antemortem* (J. Wesp, 2014).

HSJN240 (Supplementary Figure 11) is also a middle adult, morphologically male (coherent with inferred genetic sex) (Supplementary Table 4), with moderate amounts of biomechanical stress on his upper limb and early signs of osteoarthritic changes on his lower lumbar vertebrae. He has light amounts of dental wear and dental calculus; however, three maxillary teeth were lost *antemortem* and another molar has a severe cavity that would have likely resulted in another tooth loss (J. Wesp, 2014).

HSJNC81 (Supplementary Figure 11) is a young adult, morphologically male (coherent with inferred genetic sex) (Supplementary Table 4), between 20–34 years old with a light amount of dental wear and moderate amounts of dental calculus, with no major dental pathologies (J. Wesp, 2014). Unfortunately, further bio-archeological analysis is limited due to poor preservation of the cranium and the lack of other postcranial skeletal elements. The crania exhibit light to moderate amounts of porotic hyperostosis on the cranial vault and light cribra orbitalia in the eye sockets.

2. DNA extraction and NGS libraries construction

Teeth from individuals belonging to the HSJN (n=21) and COY (n=5) (Supplementary Table 1) were selected for DNA extraction, based on their morphological indicators that associate them to an African origin (Hernández-Lopez & Negrete, 2012; Karam-Tapia, 2012; Meza, 2013; Ruíz-Albarrán, 2012), or due to their state of conservation, respectively. Teeth were carefully cleaned with NaClO (70%) and ethanol (70%) superficially and later exposed to UV light for 1.5 minutes. The teeth were sectioned from the crown and fragmented by mechanical pressure inside the Human Paleogenomics Laboratory, a clean room with the necessary conditions to work with aDNA, at the International Laboratory for Human

Genome Research (Querétaro, Mexico). Approximately 200 mg of teeth samples were subjected to DNA extraction using previously reported protocols (Dabney et al., 2013; Rohland & Hofreiter, 2007) with minor modifications. A blank extraction control per batch was used to identify the presence of environmental and cross-sample contamination.

NGS libraries were constructed using 30 µl of DNA extract as previously reported with different indexes per sample (Meyer & Kircher, 2010) where a library construction blank was added per batch of DNA extract (NGS controls). All libraries and NGS controls were quantified with Bioanalyzer 2100 (Agilent) to be pooled equimolarly and sequenced in an Illumina NextSeq550 (2x75 middle output) where 217,912,556 and 205,641 reads were obtained in total for the shotgun library samples and NGS controls, respectively (Supplementary Figure 2).

3. NGS quality control and human mapping

Obtained reads (R1 and R2) from each sample were merged (at least 11bp of overlap between pairs) and adapter trimmed with AdapterRemoval 1.5.4 (Schubert et al., 2016) with a quality filter >30, and minimum length 30 bp. The resulting merged reads were used to map to the human genome (hg19) using BWA 0.7.13 (aln Algorithm) (Heng Li & Durbin, 2009), unmapped reads were used for metagenomic analysis with MALT 0.4.0 (Vågene et al., 2018).

4. MALT metagenomic analysis

Using the NCBI ftp server the whole viral NCBI Refseq database was downloaded on February 2018 containing 7530 viral genomes, including human pathogens. The Viral database was formatted with malt-build (default parameters) to create an index that was used in all our analyses without any modification.

The unmapped (non-human) reads were aligned with MALT 0.4.0 using blastn and SemiGlobal mode with a minimum 85% of identity (--minPercentIdentity) and a e-value of 0.001 (--e), the remaining parameters were used as default and resulting files were analyzed in MEGAN 6.8.0 (Huson et al., 2016).

5. In-solution enrichment assay design

Eight viral families were included in our capture design, which comprised 25 complete publicly available reference genomes, 3 consensus genomes (HBV, Bocavirus, Lentivirus), a selection of genes (*Herpesviridae*) and a consensus sequence from a selection of genes (VARV), as shown in Supplementary Table 2.
The HBV majority consensus genome (>50% conservation per site) was constructed using an alignment of modern references (A-H genotype) and a well-covered (>5x coverage) ancient genotype (Mühlemann, Jones, et al., 2018) (LT992459).

Thirty VARV genes were chosen for a consensus sequence construction based on three categories; replication (J6R, A24R, A29L, E4L, A50R, A5R, D7R, H4L, E9L), structural (A27L, A25, D8L, H3L, L1R, A33R, B5R, A16L), and immune host regulation (B18R, A46R, B15R, K7R, N1L, M2L, E3L, H1L, B8R, D9R, D10, K3L) that were obtained from all the available VARV genomes including three ancient genomes (NCBI GenBank 2019) (Duggan et al., 2016; Pajer et al., 2017). The resulted genes were aligned in AliView (MUSCLE algorithm)(Edgar, 2004; Larsson, 2014) to generate a majority consensus for every gene. The generated consensus sequences targeted <20% of the VARV whole genome.

For the *Herpesviridae* family a total of 19 genes where selected, six from Herpes Simplex Virus 1 (UL30, UL31, UL19, UL27, US6, UL10), six from Human Cytomegalovirus (UL54, UL53, UL86, UL115, UL75, UL83), and seven from Epstein Bar Virus (ORF9, ORF69, ORF25, ORF47, ORF8, vIRF2, K5). GenBank IDs are shown in Supplementary Table 2.

Selected genes from VARV and *Herpesviridae* were defined as 40 bp or 60 bp upstream the start codon, and downstream the stop codon, respectively, in order to ensure a uniform coverage of the entire coding region in case of a positive sample.

The selected sequences were further analyzed by Arbor Biosciences (Ann Arbor, MI, USA) to remove low complexity regions and sequences similar (through blastn) to the human genome (hg38), which resulted in 19,147 ssRNA different probes of 80 nt in length targeting the customized regions each 20 nt.

6. In-solution enrichment

The capture assay was performed following the manufacturer's protocol (version 4.01) and using between 30-90 ng (depending on the availability) of the indexed libraries; probes were hybridized with aDNA libraries at 60ºC during 48h.

In order to obtain whole genomes, one or two independent rounds of capture assays were performed for the B19V-positive samples (i.e. the same library was captured twice, but not consecutively). The HBV-positive library was only captured once. To evaluate the enrichment on a library with no hits to any of the eight viral families (except for a false positive human Alu sequence with homology to Poxviruses), the library from individual HSJN177 was used as negative control for the capture assay (capture control).

qPCR was carried out using primers for the adaptors of each post-capture library to estimate the number of cycles needed for amplification before reaching the plateau phase to avoid high frequency of PCR clones. Amplified libraries were then purified following the SPRISelect Magnetic Beads manufacturer's protocol (Beckman Coulter), and quantified with the Bioanalyzer 2100 (Agilent). The pooled libraries were sequenced (Supplementary Figure 1) to obtain entire HBV (Figure 2c) or B19V (Figure 2a) genomes on an Illumina NextSeq550 (2x75 middle output).

7. Metagenomic comparison between pre- and post-capture sequencing

NGS reads were analyzed in the same way as the shotgun data (Supplementary Methods 3) in order to obtain non-human reads from each capture experiment, which were analyzed with MALT 0.4.0 (Vågene et al., 2018) (Supplementary Methods 4). The resulting viral abundancies were normalized (default parameters) and compared with MEGAN 6.8.0 (Huson et al., 2016) to assess the capture efficiency on the targeted viruses (Figure 1c).

8. Viral datasets

HBV/DS1:
For BWA analysis 38 genomes were included in total. GenBank IDs:

Genotype A: HE974381, HE974383, AY934764, GQ331046

Genotype B: B602818, AB033555, AB073835, AB287316, AB241117

Genotype C: AB111946, X75656, AB048704, AF241411, AP011102, AP011106, AP011108, AB644287

Genotype D: FJ899792, GQ477453, JN688710, HE974373, FJ904430, AB033559

Genotype E: HE974384

Genotype F: AY090458, AB116654, AY090455, DQ899144, HE974369, AB116549, AF223962, AB166850

Genotype G: AP007264

Genotype H: AB516395

Genotype J: AB486012

Ancient: LT992443, LT992459

Outgroup (Woolly Monkey): AF046996

HBV/DS2:

Contains the union of curated datasets used in four previous studies (Drexler et al., 2013; Krause-Kyora et al., 2018; Mühlemann, Jones, et al., 2018; Paraskevis et al., 2015), from which only not duplicated HBV genomes were considered and resulted in a total of 593 sequences from genotypes A-J and non-human primates (Gibbon, gorilla, chimpanzee). GenBank IDs:

EU306725, HE981175, JQ027312, JQ429078, AY167091, AB300371, KU605533, AB014366, EU859930, AB365453, AY161138, FN545828, FM199978, FJ692596, AB116092, AB194951, KJ854685, KM606737, KU605537, FJ692558, JN182327, AM184125, AY233275, KF214660, KJ854693, KP168428, FN545833, KU964274, KX276846, JX026879, AB014371, JQ801498, KF922430, JQ023661, EU366129, EU859952, JN182323, KT151612, KF922415, FJ692592, FJ692608, FN545826, AY233290, JQ707397, DQ890381, AY233280, KF922434, KX357650, DQ089795, AB219533, JQ429081, KP168435, AB111120, AP011099, KJ803826, KT364751, KU679947, KR013871, AB049610, HQ700576, JQ027317, JX507211, KJ410494, GQ358158, GQ924642, HM011488, KU964236, DQ089790, KC774297, EU498227, KJ410496, KX276836, X75665, EU939568, KF214673, KF873541, KJ803766, KJ803823, GQ475321, HQ700564, KF873514, KJ803809, HM011479, AB367420, KF873536, AB670295, EU796069, EU939539, KX276850, AB176642, AP011106, EU670263, KU964045, KX276848, KC774238, AB697502, AB900109, KJ803818, KC774180, KC774226, GQ377586, HQ700516, HQ700575, KX276853, AB112472, DQ089768, DQ089802, JN827415, KC774214, KR013859, AB116654, AB214516, AY090458, DQ899142, DQ899145, DQ899148, KC774444, AB112063, AB195931, JQ040132, KF214670, KJ803790, HQ700522, KC774196, AB670259, AB697510, AY206386, D23682, X75658, X75663, KF199901, AB059659, AB059660, JN792921, JQ272888, KJ676694, KJ638660, KJ638662, JN688720, HQ700456, JF828925, AB900113, DQ089777, AB300361, AB367392, DQ089804, FJ386626, FJ386644, KX276841, AB113878, KX276855, AB198079, JX870000, KR013837, KJ803777, KJ803779, KU679937,

KU679951, KC774244, JQ027324, AB115417, AB195930, AB931170, KF873526, EU939597, FJ899767, GQ358157, GQ924643, KX276800, AB073821, AB287315, AB713528, AY800392, DQ463791, EF494381, EU939678, FJ386582, KP148582, JQ707737, KX276792, AB010289, AB073853, AB212625, AB828708, KM875420, KP148452, KP406278, JN827419, GQ358137, GQ358151, FJ032342, GQ924645, KX276791, KX276795, KX276825, KX276827, KJ803796, KP148414, KJ173297, DQ995804, EU522072, GQ358146, GQ358148, GQ924624, KX276772, KX276798, KX276806, AB900098, HM011483, KJ173401, JQ027311, JQ027330, KC774370, GQ924626, AB287323, DQ993684, DQ995802, KX276786, KX276815, GQ924637, HM011476, KU964383, KX276796, KX276819, JX661471, KJ803805, HM011466, KJ790200, KX276787, AB900107, FJ023631, HM011499, HM011503, JQ027334, KX276783, KX276785, KX276794, KX276830, AB231909, AB642101, AB931169, KX276817, AB670298, HM011487, JQ027313, KX276844, KU964358, AB670263, AB246340, EU939634, FJ562260, KX276770, KX276812, GQ924630, HM011478, KJ803795, FJ386601, DQ089785, JQ040167, JN827421, HM011495, GQ377632, FJ787452, DQ089788, AY641559, AB670258, KJ410515, KC774351, KC774302, KC774240, KC774357, KC774352, HM011482, KJ803820, KP659250, GQ924621, FJ386648, GQ924641, KX276807, KX276821, AB031267, AB073849, AB106884, AB219429, HM011471, HM011490, AB555498, AB642093, EU158262, AB073836, AY206383, KJ803817, KP659249, HQ700546, HM011496, KJ803808, AY206391, GQ924656, EU939677, KX276813, AB100695, AB241117, KX276797, KJ173379, KJ410502, JF436921, HM011475, KX276858, KJ173342, AB274977, HM363586, HM363592, HM363603, EU239220, HM363611, KU736913, AB106564, FN594760, GQ161775, HM011467, AB074047, FN594751, FN545823, DQ315778, FJ899783, KX276774, AY641563, AF297621, HQ700506, GQ924635, HQ700517, KF873544, AB670285, EU522071, DQ399006, GQ184322, JF754588, GU456678, GQ167302, JN257177, JN688713, L27106, HQ700458,

HQ700449, GQ922002, EU414139, JN664913, JN642163, JN642159, JN257190, JN257162, JN040768, JN664922, JQ707699, JN688695, JN688712, KU736927, KJ647355, JN664919, KF679990, KF192832, KM524358, KC875342, JN688683, JN040769, JF754617, HQ700510, GU456658, GQ477452, FJ904427, JN642135, KU668435, EU594434, EU594432, EU414140, AY090452, AJ627220, GU456674, GQ477456, FJ904438, FJ904395, JN664931, JF754611, EU155893, DQ464174, DQ304548, JX470760, JN792912, JN664936, JN257202, JF754592, KP168419, AJ627218, AB674428, AB555500, AB330367, GU456679, GQ205385, GQ205382, GQ205380, JN040762, JF754612, GU456643, JN040822, AB674416, AB270541, X80926, GQ922005, FJ904445, GU456651, FJ904439, JN664932, JN642136, JN642133, JQ707529, AB188241, AB048701, X65258, GU456665, GQ922003, GQ477453, FJ904402, EU414141, AJ627222, AJ344116, JN257165, JN257160, JN040779, JF754631, AF043594, AB674425, KX357622, KM524338, KJ843187, GU456684, JN688710, JN664921, JN257172, JN040818, KM577668, KJ470893, FJ904433, FJ904422, DQ464173, AB674424, AB555496, AB119255, JF754597, HQ700513, GU456682, GU456669, GU456654, GU456637, FJ904426, JN688711, JN664920, JN642149, JN040766, KP090181, KJ647353, DQ486025, AB674427, AB823660, AB823659, AY330914, AY077735, EU155821, AY781187, AY781186, AY781182, AY330917, AY330916, AY330915, AY330913, AY330912, AY330911, AY077736, AF242586, AB823662, AB823661, AB823658, JQ664503, EU155829, AJ131571, AJ131574, AJ131569, FJ798098, AB032432, AF222322, AF498266, AF193864, AF193863, AM117396, MG585269, AB076679, AB116084, AB453988, AY738142, GQ477499, AY934764, FJ692556, FJ692598, FJ692611, GQ161813, GQ331046, AB073858, AB033555, AB219429, AB219430, AP011089, AB073835, AB287316, AB287318, AB287320, AB287321, DQ463789, DQ463792, AB241117, DQ993686, AB111946, AB112066, AB112472, DQ089767, X75656, X75665, AB048704, AB048705, AF241411, AP011100, AP011102,

AP011103, AP011106, AP011108, FJ899792, JN642140, GQ477453, GQ477455, JN642160, JN642163, JN688710, JN688711, GQ922005, HE974378, KJ470893, KJ470896, KJ470898, FJ904430, FJ904436, AB033559, AB048701, AB048702, AB188243, AB210818, AM494716, AY796031, AY902768, DQ315779, X80925, X75657, X75664, AY090458, AB116654, FJ657525, AY090455, AY311369, DQ899144, DQ899146, AB116549, X75663, AF223962, AB166850, AB056513, AB064312, AB059660, AB375163, AY090454, AY090457, AB486012, AB032433, AB493845, DQ298161, DQ336682, GQ205385, GQ331047, GQ475343, GQ922004, JN040772, KF679991, KJ470892, AY330911, AJ131571, AJ131567, AF193863.

1 Ancient HBV genome from a Korean mummy (Kahila Bar-Gal et al., 2012): JN315779

1 Ancient HBV genome from a Naples mummy (Patterson Ross et al., 2018): MG585269

3 Ancient HBV genomes from Neolithic Germany period (provided kindly from the authors) (Krause-Kyora et al., 2018): Petersberg, Sorsum, Karsdorf.

12 Ancient HBV genomes from Bronze to Medieval Eurasian period (Mühlemann, Jones, et al., 2018): LT992439, LT992442, LT992444, LT992440, LT992441, LT992454, LT992455, LT992459, LT992448, LT992447, LT992443, LT992438

1 Ancient HBV genome from Egyptian Mummy (provided kindly from the authors) (Neukamm et al., 2020): Abusir1543

1 Ancient HBV genome from an African individual in the HSJN (provided kindly from the authors) (Barquera et al., 2020): HSJN150

1 Ancient HBV genome from the present study: HSJN194

HBV/DS2.1

56 genomes assigned to Genotype A, based on our ML analysis plus the ancient genome from the present study (HSJN194)

B19V/DS1:

For BWA and BLASTN analysis 14 genomes were included. GenBank IDs:

13 B19V genotype 1 to 3: KT268312, AY504945, FJ591158, EF216869, AY064476, DQ333427, AB550331, AY582124, DQ408305, FJ265736, AJ249437, NC_004295, NC_000883.

1 Outgroup (Bovine Parvovirus): NC_001540.

B19V/DS2:

All B19V genomes retrieved in the NCBI database were downloaded using the next search command "*human parvovirus b19[organism] not rna[title] not clone[title] not clonal[title] not patent[title] not recombinant[title] not recombination[title] and 3000:6000[sequence length*]" which considers only whole genomes (3-6kb resulting in a total of 109 B19V genomes. GenBank IDs:

Genotype 1 to 3:

MH151117, MH201455, MH201456, HQ340601, NC_000883, NC_000883, Z70528, AB030673, AB030693, AB030694, AB126262, AB126263, AB126264, AB126265, AB126266, AB126267, AB126268, AB126269, AB126270, AB126271, AB550331, AF113323, AF162273, AJ249437, AJ717293, AJ781038, AY028237, AY044266, AY064475, AY064476, AY083234, AY386330, AY504945, AY582124, AY582125, AY647977, AY903437, DQ225148, DQ225149, DQ225150, DQ225151, DQ234769, DQ234771, DQ234775, DQ234778, DQ234779, DQ293995, DQ333426, DQ333427, DQ333428, DQ357064, DQ357065, DQ408301, DQ408302, DQ408303, DQ408304, DQ408305, EF216869, FJ265736, FJ591158, FN598217, FN598218, HQ340602, KC013305, KC013308, KC013312, KC013313, KC013314, KC013316, KC013321, KC013322, KC013324,

KC013325, KC013327, KC013329, KC013331, KC013332, KC013333, KC013338, KC013340, KC013343, KC013344, KC013346, KC013351, KF724386, KF724387, KM065414, KM065415, KM393163, KM393164, KM393165, KM393166, KM393167, KM393168, KM393169, KR005640, KR005641, KR005642, KR005643, KR005644, KT268312, KT310174, KX752821, M13178, M24682, NC000883, Z68146, Z70560, Z70599.

10 ancient B19V coding regions from genotype 1 and 2 (Mühlemann, Margaryan, et al., 2018) (https://github.com/acorg/parvo-2018/tree/master/consensuses):

DA251-user, DA336, DA337n, DA66, NEO105, RISE569, VK143, VK154, VK477, VK6.

3 Ancient B19V coding regions from the present study:

HSJNC81, HSJN240, COYC4

9. Ancient viral genomes reconstruction and authentication

After enrichment, the non-human (unmapped) data was used to generate HBV or B19V consensus genomes as follows.

HBV: Reads were competitively mapped to HBV/DS1 with BWA (aln algorithm) (Heng Li & Durbin, 2009) with seedling disabled (-l 1050)(Schubert et al., 2012). The reference with the most hits was used to map individually (BWA, same parameters as above) and duplicates were removed with Samtools rmdup (H. Li et al., 2009).

B19V: Ancient B19V genomes were reconstructed as previously reported (Mühlemann, Margaryan, et al., 2018). Using a pipeline that includes a first BWA competitive mapping (seedling disabled). If >50% of the B19V genome was covered, samples were considered positive and queried with blastn (-evalue 0.001) to B19V/DS1. To avoid local alignments, only global hits (>85% of read) to B19V were conserved. To control for a likely false-positive taxonomic assignation due to the biased viral database, filtered reads (HBV and B19V) were subsequently analyzed with megaBLAST (default online parameters) using the whole NCBI non redundant (nr) database. Only reads for which the best hit was to HBV or B19V were kept for downstream analysis (Supplementary Table 3, Supplementary Figure 1). Filtered data (Supplementary Table 3) was mapped to a reference dataset including sequences representative of genotypes 1, 2, 3 in GeneiousPrime 2019.0.4 (Kearse et al., 2012) (medium sensitive/fast, 5 iterations) to choose the reference with the largest coverage and duplicates were removed.

The alignments were analyzed with mapDamage 2.0 (Jónsson et al., 2013) to generate damage patterns (Supplementary Figure 3) and to rescale the base quality scores of the likely damaged bases (--rescale option). The quality-rescaled alignments were used to generate a consensus genome with Bedtools 2.21.1 (bamtofastq) (Quinlan & Hall, 2010).

10. Phylogenetic analysis

HBV/DS2 and B19V/DS2 were aligned independently in Aliview (Larsson, 2014) (Muscle algorithm (Edgar, 2004)) and curated manually to have the same lengths. The alignments were evaluated in jModelTest 2.1.10 (Darriba et al., 2012) using a corrected Akaike (AICc) and Bayesian (BICc) Information Criterion tests that supported with 100% confidence the evolutionary models used in our maximum likelihood analysis in RAxML (Stamatakis, 2014).
To test the temporal structure of our ML trees, a root-tip dated analysis was performed on Tempest 1.5.3 (Rambaut et al., 2016) for both DS2 (B19V, HBV) in

the presence or absence of ancient sequences, and without the sequences presented in this study (Supplementary Figure 6 and 7).

In the case of HBV, a more specific analysis was performed only on the Genotype A to find a higher temporal, in the presence or absence of ancient sequences, and without the HSJN194 HBV genome presented in this study (Supplementary Figure 6).

For the B19V/DS2 the temporal structured suggested by root-tip distances analysis was corroborated using a date randomization test (DRT) with TipDatingBeast 1.0.5 (Rieux & Khatchikian, 2017) and BEAST 2.5.1 (Drummond et al., 2012) (Supplementary Figure 7). Since the DRT and the root-tip dated analysis suggested a temporal structure for the B19V/DS2, a coalescent dated tree was generated in BEAST 2.5.1 (Drummond et al., 2012) for B19V using a relaxed and strict clock; both with different priors (coalescent constant, exponential and Bayesian skyline population priors), with an *a priori* substitution rate interval of $1x10^{-3}$-$1x10^{-7}$ s/s/y (Mühlemann, Margaryan, et al., 2018). For the Colonial genomes used in this study a uniform sampling was indicated using the radiocarbon dates for the individual HSJN240 (495 ± 166 ybp). When radiocarbon dating was not possible an archeological date interval was set for HSJNC81 (332.5 ± 269 ybp) and COYC4 (320 ± 400 ybp), based on the archeological estimates of both sites.

The strict molecular clock analyses were performed with a 50 million MCMC sampled each 5000 generations, while the relaxed molecular clock with exponential population was run with a 250 million MCMC sampled each 5000 generations, and the relaxed molecular clock with coalescent constant and Bayesian Skyline population priors were ran with 250 million MCMC and with 350 million MCMC sampled each 5000 generations. Both files were mixed with a 25% burn in LogCombiner (Drummond et al., 2012).

All the Bayesian analyses were mixed and reached convergence (>200 ESS) as estimated in Tracer 1.7 (Rambaut et al., 2018) (Supplementary Table 5). The first

25% of the generated trees where discarded (burn in) and a Maximum Clade Credibility Tree with median ages was created with TreeAnnotator (Drummond et al., 2012) (Supplementary Figure 5).

11. Radiocarbon dating

Radiocarbon ages were determined at the Physics Institute of the National Autonomous University of Mexico (UNAM) for the individuals in this study with complete skeletons (HSJN194 and HSJN240). From these individuals, phalange bones (left hand) were cleaned, dried and powdered to be digested on a HCl 0.5M solution followed by a NaOH 0.01M and HCl 0.2M treatment. Collagen was then filtered (>30 KDa) and graphitization was performed on an AGEIII (Ion Plus). $^{14}C$, $^{13}C$ and $^{12}C$ isotopes were analyzed from graphite in a Tandetrón (High Voltage Engineering Europa) mass spectrometer with a 1 V energy accelerator. Radiocarbon dates were estimated based on InCal13(Reimer et al., 2013) calibration curve and corrected with OxCal v4.2.4. (Bronk Ramsey, 2013)

12. Sr isotope analysis

Tooth enamel was carefully extracted with the aid of dentist tools. The material underwent several cleaning procedures before crushing to a 50 µm grain size with an agate mortar. Chemicals used for this purpose included 30 % $H_2O_2$ and 1-1.5 N $HNO_3$. In between, deionized water (Milli-Q) rinses were performed. Ultrasonic bath (USB) was used to accelerate processes. After obtaining the desired grain size, samples were treated with 30 % $H_2O_2$, 1N $NH_4Cl$ and alternated water washes.

In order to get rid of any secondary contaminant or any post-mortem external agent that could alter the Sr isotopic values, tooth samples were treated with a three-step leaching technique: the first leachate is obtained with 0.1N acetic acid for 30 minutes (USB). The solution is decanted and dried under infrared light (Lix 1). The residue was leached for 15 minutes in 1N acetic acid (USB) and subsequently stored overnight for 12 hours in the same acid. The solution was decanted and dried to obtain the second leachate (Lix 2). The residue (Res) is dissolved in 8N $HNO_3$ as well as Enamel Lix 1 and Enamel Lix 2 in closed Teflon® beakers on a hot plate at 90°C. A total of three aliquots from each molar were obtained from this leaching process. The efficiency of this procedure on human teeth has been previously validated (Solís-Pichardo et al., 2017). In this work, the expected outcome of the leaching process is shown in Supplementary Figure 9 and Supplementary Figure Table 6; whereas in individual HSJN240 there is not much variation between the three $^{87}Sr/^{86}Sr$ leachate aliquots, sample HSJN81 displays significant differences, and demonstrates the elimination of secondary Sr contamination during the leaching procedure. After sample digestion, Sr from teeth and bone samples was extracted with Sr-Spec (EICHROM®) ion exchange column chemistry. Detailed analytical procedures are described in (Solís-Pichardo et al., 2017).

Sr isotope analysis was carried out with a Triton Plus (Thermo Scientific) thermal ionization mass spectrometer with 9 Faraday collectors at the "Laboratorio Universitario de Geoquímica Isotópica" (LUGIS, UNAM). Sr was measured as metallic ions with 60 isotopic ratios that were normalized for mass fractionation to $^{86}Sr/^{88}Sr$ =0.1194. The mean value for the NBS 987 Sr standard was $^{87}Sr/^{86}Sr$ = 0.710254±0.000012 (±1$sd_{abs}$, n=86) and the analytical blank yielded 0.23 ng Sr.

$^{87}Sr/^{86}Sr$ ratios were performed on the enamel teeth (crowns) of the individuals HSJNC81 and HSJN240.$^{87}Sr/^{86}Sr$ in teeth enamel from adults is formed uniquely during the development of a human being (approx. first 5 years of life) and reflects the geographic location at birth (Bentley, 2006). Similar analyses were done on HSJNC81 and HSJN240 using their parietal and phalange bone, respectively. In contrast to tooth enamel, bone $^{87}Sr/^{86}Sr$ remodels over an individual's existence and indicates the place where the person spent his or her last years of life (Solís-

Pichardo et al., 2017). Diagenetic and other secondary alteration processes in bone may change the Sr isotopic values, but it was demonstrated that these variations are within a narrow range of approximately 6% (Solís-Pichardo et al., 2017). This was achieved by comparing 17 soil samples from the Mexican Altiplano (rocks of the central Trans Mexican Volcanic Belt (TMVB)), which yielded a $^{87}Sr/^{86}Sr$ mean of 0.70449 ±0.00025 (1 sd), with the $^{87}Sr/^{86}Sr$ bone mean of 0.70478 of 27 *Teopancazco* (Teotihuacan) individuals (Solís-Pichardo et al., 2017). In the case of the two individuals analyzed in this study, bone $^{87}Sr/^{86}Sr$ values 0.70672 (HSJNC81) and 0.70755 (HSJN240) (Supplementary Table 6), are comparable to those obtained from soil samples from the eastern TMVB rim in Veracruz with a mean $^{87}Sr/^{86}Sr$ of 0.70703 (n=6) (Solís-Pichardo et al., 2017). For West African igneous and metamorphic rocks, a mean value $^{87}Sr/^{86}Sr$ of 0.71044 was obtained (n=20, Supplementary Figure 9). Data are compiled in Supplementary Table 7 with their corresponding references.

**<u>Supplementary Discussion</u>**:

1. Preservation of ssDNA in viral genomes

B19V has a broad tropism and infects precursors of the erythroid lineage (Broliden et al., 2006), however DNA has been recovered from other few non-erythroid lineage cells that support B19V infection with abortive replication (Luo & Qiu, 2015). Thus it is possible to find it from teeth roots that are vascularized during an individual's life (Key et al., 2017).

At the same time, the presence of ssDNA genomes as B19V in previous studies (Mühlemann, Margaryan, et al., 2018) and in our samples was unexpected since the NGS libraries were constructed for dsDNA as a template. Not to mention that ssDNA is more prone to depurination and deamination than dsDNA reducing its stability through time (Lindahl, 1993). Nevertheless, dsDNA intermediates have described to form during the virus replication cycle (Ganaie & Qiu, 2018). Throughout viral infection, the replicating genomes are present at different steps allowing the construction of NGS libraries on these cell-free dsDNA intermediates. Furthermore, the ITRs of the B19V genome (Figure 2b) are in a hairpin-like dsDNA conformation inside circulating virions and throughout the majority of the Parvovirus replication cycle (Luo & Qiu, 2015), and are less prone to degradation than ssDNA (Lindahl, 1993).

Conversely, HBV has a partial dsDNA genome where the negative-strand is completely circular with a ssDNA region (Yuen et al., 2018), which forms dsDNA at intermediate steps during its replication cycle (cccDNA) (Grimm et al., 2011). This ssDNA region has low coverage in our ancient HBV genome (Figure 2c), as well as in previous HBV ancient genomes (Krause-Kyora et al., 2018; Neukamm et al., 2020), consistent with a higher ssDNA degradation over dsDNA (Lindahl, 1993); although, we cannot discard the possibility that the recovered reads were obtained from cell-free dsDNA intermediates present at low frequency.

HBV (Yuen et al., 2018) and B19V (Janovitz et al., 2017) are also capable of integrating into the human host genome, which could explain the discrepancy of mutation rates among modern and ancient strains(Simmonds et al., 2019). B19V is able to integrate in peripheral blood stem cells (CD34+) (Janovitz et al., 2017), but the poorly covered ssDNA region does not support its integration as dsDNA into the human genome. Similarly the unusual coverage found at the ssDNA region in HBV from previous studies (Krause-Kyora et al., 2018) and ours does not support that we are identifying integrated DNA, not to mention that HBV integrates mainly in hepatocytes (Furuta et al., 2018). Further analyses are needed to formally discard the possibility that the recovered ancient viral genomes to date were integration events after an HBV or B19V infection, and instead elucidate if the aDNA retrieved is coming from circulating virions or from replication intermediates found as cell-free DNA (Cheng et al., 2019). Additionally, clarifying the nature of the aDNA would let us understand better the mechanism by which ssDNA from viruses is conserved throughout time.

2. B19V Infections during the Colonial period in Mexico

B19V is part of the Erythoparvovirus genus and infects precursors of the erythroid lineage. It has a broad tropism and has been recovered from multiple human tissues such as bone marrow, liver, heart, placenta (Luo & Qiu, 2015), tonsil, testicles, kidney, muscle, thyroid, brain (Pyöriä et al., 2017) and bones (Mühlemann, Margaryan, et al., 2018; Toppinen et al., 2015). B19V is the causative agent of fifth disease (*erythema infectiosum*) common in children, who present symptoms such as fever and rashes (Broliden et al., 2006).

In several reports (Anderson et al., 1985; Davidkin et al., 1998; De Los Ángeles Ribas et al., 2019; Rezaei et al., 2016), children and young adults with measles-like symptoms have been negative to measles, and instead tested positive to B19V (Anderson et al., 1985; Davidkin et al., 1998; De Los Ángeles Ribas et al., 2019; Rezaei et al., 2016) or rubella (Anderson et al., 1985; Davidkin et al., 1998; De Los Ángeles Ribas et al., 2019; Rezaei et al., 2016), which produce a similar kind of rash and fever. There are historical records that describe the treatment of measles outbreaks at the HSJN in 1531 (Meza, 2013), nevertheless in the absence of immunological and molecular tools, the accurate diagnosis of measles during the Colonial period of Mexico City was impossible and possibly misinterpreted B19V or rubella infection, as has been demonstrated in modern cases. Our study does not exclude the notorious role of measles during the Colonial outbreaks (as strongly suggested by historical records), but provides evidence of the presence of B19V during the Colonial period of Mexico City, raising a debate about the paradigmatic etiology of the supposed measles epidemics reported in historical records (Malvido & Viesca, 1982; Mandujano-Sánchez et al., 1982; Somolinos d'Árdois, 1982). Further developments in aDNA methodologies are needed to have an unbiased screening of RNA (measles, rubella, influenza) and DNA viruses simultaneously, as carried out in recent studies (Düx et al., 2020).

Furthermore, B19V is also associated with liver failure, fulminant hepatitis, arthralgias and severe anemias (aplastic crisis) in adults (Broliden et al., 2006; Chisaka et al., 2003). The anemia caused by B19V is explained by an abrupt cessation of erythroid precursors cells in the bone marrow (Chisaka et al., 2003). The HSJN skeletal collection has a notably higher rate of cribra orbitalia and porotic hyperostosis compared to other Colonial archeological sites (Castillo-Chavez, 2000). These skeletal indicators are caused by an irregular hematopoiesis in the bone marrow and are typically associated with genetic anemias such as thalassemia and sickle cell anemia (Angel, 1966), as well as to nutritional stress or parasitic infections (Walker et al., 2009). From the individuals with ancient B19V genomes in our study, the individual HSJNC81 presented cribra orbitalia in the eye sockets and porotic hyperostosis on the cranial vault. The B19V ancient genome from the HSJNC81 is highly similar (Figure 3b, Supplementary Figure 5) to the strain V9 (AJ249437) recovered from an infant with severe anemia and G6PD deficiency (Nguyen et al., 1999). Previously, these frequent anemia-associated cranial marks in the HSJN were proposed to be caused by an unknown infectious disease (Castillo-Chavez, 2000), however it might be possible that these marks were a consequence of severe anemia caused by the ancient B19V presence in individual HSJNC81 or enhanced in the presence of other hematological disorder (Broliden et al., 2006).

Our study cannot discard a genetic cause for the osteological indicators of anemia on the HSJNC81, since the loci for thalassemia, sickle cell anemia and G6PD deficiency were not covered with our human-mapped NGS data. Nevertheless, B19V infection is strongly enhanced (transient aplastic crisis) with other hematological disorders, such as thalassemia, sickle cell anemia, nutritional stress, malaria and iron deficiency, which become severe and even lethal when present together (Broliden et al., 2006). In predisposed individuals 70-80% aplastic episodes are caused by B19V infection (Heegaard & Brown, 2002).

The infection with B19V results generally in a lifelong immunity; nevertheless, in immunosuppressed and also some healthy individuals with an inability to develop a neutralizing immune response, virus infection lead to chronic anemia or arthropathies (Heegaard & Brown, 2002).

The presence of B19V in the Americas (non continental) before the Spanish arrival has been reported previously for Greenland (genotype 2 in Viking burial) (Mühlemann, Margaryan, et al., 2018). Additional analyses on Pre-Hispanic remains are needed to explore the presence of this and other viral genotypes in Mexico to further understand the epidemiology of the Colonial population.

3. Hepatitis during the Colonial Period of Mexico

The HSJN was probably the most important hospital for the Indigenous inhabitants during the Colonial period (Meza, 2013), though patients also included individuals of African or mixed heritage, and they treated notorious epidemics, such as the *Cocoliztli* of 1576. The autopsies on the dead bodies from the *Cocoliztli* of 1576 were performed at the HSJN by Dr. Francisco Hernandez ("Proto-Medico" of the New Spain and former personal physician of King Phillip II of Spain) and Dr. Alfonso de Hinojoso (physician of the HSJN) (Acuña-Soto et al., 2004; Marr & Kiracofe, 2000; Somolinos d'Árdois, 1982).

Dr. Hernandez described this epidemic as "contagious lethal fevers, the urine was green and black, the tongue was dry and black, blood emanated from ears and nose, the pulse was fast and weak, the eyes and the whole body were yellow and followed by delirium, heartache and great anxiety" (Acuña-Soto et al., 2004; Marr & Kiracofe, 2000; Somolinos d'Árdois, 1982). At autopsy, the liver was greatly enlarged, the heart was black with a yellow liquid and black blood, the spleen and lungs were black, and the rest of the body was pale (Acuña-Soto et al., 2000; Malvido & Viesca, 1982; Somolinos d'Árdois, 1982).

The disease affected mainly the Indigenous inhabitants, followed by admixed populations (Indigenous with Spaniards), later Africans and ultimately the Spaniards (Acuña-Soto et al., 2000; Malvido & Viesca, 1982; Somolinos d'Árdois, 1982). Independently, Dr. Hinojoso wrote a report based on the autopsies of the same epidemic where he described that the liver was extremely enlarged and hard, as well as splenomegaly (Acuña-Soto et al., 2000).

Different interpretations of the etiology of this epidemic have been done and include yellow fever, plague, influenza, leptospirosis, hepatitis, malaria, typhus, hemorrhagic fever, smallpox, measles, typhus (Acuña-Soto et al., 2000; Malvido & Viesca, 1982; Marr & Kiracofe, 2000) or a synergic effect of the coinfection with different pathogens (Somolinos d'Árdois, 1982), and more recently the infection of *Salmonella enterica* (*Cocoliztli* 1545) based on an aDNA study (Vågene et al., 2018). Although some of these diseases and pathogens fit with many of the historically described symptoms none of them independently entirely satisfies the complex characteristics of the 1576 *Cocoliztli*.

Marr and Kiracofe (Marr & Kiracofe, 2000) interpreted the color of the blood and body reported in these autopsies as a symptom of a severe intravascular hemolysis cause by the metabolization of the hemoglobin into bilirubin, which, in excess, produces a yellow color in the eyes and body as well as a change of color of the urine, attributed to arthropod borne pathogens (malaria, yellow fever virus, and dengue virus) (Marr & Kiracofe, 2000). On the other hand, Acuña-Soto *et al.* (Acuña-Soto et al., 2004) interpreted the symptoms as a severe liver damage that caused intense jaundice, hepatomegaly, and dark urine, which included hepatitis encephalopathy (anxiety and dementia) with portal hypertension (splenomegaly) and bleeding as seen in severe untreated liver damage (Acuña-Soto et al., 2004).

These signs are also observed in chronic HBV infections, as it proliferates in hepatocytes producing hepatitis and jaundice (Yuen et al., 2018). B19V has also been retrieved from liver and is associated with liver failure, fulminant hepatitis and jaundice (Broliden et al., 2006; Luo & Qiu, 2015).

Additionally, our radiocarbon dating for the individual HSJN194 (HBV) and HSJN240 (B19V) suggest these individuals died between 1472-1625 CE and 1442-1608 CE (years calibrated for 1σ), respectively (Supplementary Figure 10), which overlaps with the period of time when the hepatitis symptoms where reported after the 1576 *Cocoliztli* at the HSJN (Acuña-Soto et al., 2004; Marr & Kiracofe, 2000; Somolinos d'Árdois, 1982). Our data, however, is not sufficient to elucidate the age when the individual HSJN194 acquired HBV, if it was vertically or horizontally transmitted, nor if he presented an acute or chronic infection in any way related to his cause of death. Nevertheless, the presence of hepatotropic viruses as HBV and B19V during the Colonial period of Mexico City opens new discussions surrounding the etiology of the liver damage reports during epidemics in the HSJN.

The modern current mortality rate of both viruses is by far lower than that experienced during the *Cocoliztli* epidemics, and probably they do not represent a unique etiologic agent of this disease, instead our data could fit a scenario of synergic coinfection by different pathogens.

**Supplementary Tables:**

| Viral Family | Virus | Sequence | Reference | Length (kb) |
|---|---|---|---|---|
| *Poxviridae* | Variola virus | 30 genes | Consensus | 36.44 |
| *Hepadnaviridae* | Hepatitis b virus | Genome | Consensus | 3.26 |
| *Parvoviridae* | Human Adenoassociated virus 2 | Genome | NC_001401.2 | 4.68 |
| | Bocaviruses | Genome | Consensus | 5.54 |
| | Human Parvovirus B19 | Genome | NC_000883.2 | 5.6 |
| | Human Parvovirus B19 V9 | Genome | NC_004295.1 | 5.03 |
| *Herpesviridae* | Herpes simplex virus 1 | 6 genes | NC_001806.1 | 17.54 |
| | Cytomegalovirus | 6 genes | NC_006273.1 | 14.85 |
| | Epstein-Barr virus | 7 genes | NC_009333.1 | 27.03 |
| *Retroviridae* | Lentivirus | Genome | Consensus | 11.05 |
| | Human foamyvirus | Genome | KX087159.1 | 11.95 |
| *Papillomaviridae* | Human papilomavirus 16 | Genome | NC_001526.4 | 7.91 |
| | Human papilomavirus 5 | Genome | NC_001531.1 | 7.75 |
| | Human papilomavirus 4 | Genome | NC_001457.1 | 7.35 |
| | Human papilomavirus 1 | Genome | NC_001356.1 | 7.81 |
| | Human papilomavirus 41 | Genome | NC_001354.1 | 7.61 |
| *Polyomaviridae* | Human polyomavirus 9 | Genome | NC_015150.1 | 5.03 |
| | Human polyomavirus JC | Genome | NC_001699.1 | 5.13 |
| | Human polyomavirus BK | Genome | NC_001538.1 | 5.15 |
| | Human polyomavirus 6 | Genome | NC_014406.1 | 4.93 |
| *Circoviridae* | Cyclovirus 1 | Genome | KR902499.1 | 1.84 |
| | Cyclovirus PK5034 | Genome | GQ404845.1 | 1.78 |
| | Cyclovirus NG12 | Genome | GQ404854.1 | 1.79 |
| | Cyclovirus NG14 | Genome | GQ404855.1 | 1.79 |
| | Cyclovirus VN | Genome | KF031466.1 | 1.85 |
| | Cyclovirus VS | Genome | KC771281.1 | 1.83 |
| | Cyclovirus PK5222 | Genome | GQ404846.1 | 1.74 |

| | | | | |
|---|---|---|---|---|
| | Cyclovirus 10 | Genome | KF726984.2 | 1.79 |
| | Cyclovirus SL | Genome | KJ831064.1 | 1.71 |
| Total Length | | | | 217.77 |

**Supplementary Table 1. Viral families included in customized capture design.** Twenty-five whole genomes from different viruses were included in our customized capture design, as well as a selection of genes (Supplementary Methods 5) from *Herpesviridae* and *Poxviridae* (VARV). Raw or consensus sequences were used as described in Supplementary Methods 5.

| Virus | Sample | Shotgun hits | Capture 1 hits | Capture 2 hits | Fold Enrichment |
|---|---|---|---|---|---|
| HBV | HSJN194 | 2 | 209 | NA | 104.5x |
| | HSJN177 (negative control) | 0 | 3 | NA | - |
| B19V | HSJNC81 | 5 | 593 | 607 | 118.6-121.4x |
| | HSJN240 | 5/12* | 591 | 594 | 49.5-118.2x |
| | COYC4 | 1 | 226 | 214 | 214-226x |
| | HSJN177 (negative control) | 0 | 1 | NA | - |

**Supplementary Table 2. Enrichment yield of HBV-like or B19V-like hits.** The metagenomic analysis was carried out with MALT 0.4.0 based on the NCBI RefSeq Viral database, hits were normalized automatically between capture assay(s) and shotgun assay(s) per sample in MEGAN 6.8.0. HSJN177 was considered a negative control for capture based on the shotgun metagenomic analysis. Fold enrichment was calculated as capture hits / shotgun hits. *Two independent NGS libraries were constructed (and then captured) from two different teeth samples belonging to the same individual. NA: Not available.

| Sample | Capture | Total sequences (not human) | Mapped BWA (DS1) | Mapped Blastn DS1 (>85% identity) | BWA+Blastn | Top hit (megablast nr NCBI) | Joined reads (all capture assays) | Without duplicates | Clonality (%) | %Coverage (bp >1 read) # | Average depth# |
|---|---|---|---|---|---|---|---|---|---|---|---|
| COYC4 | 1 | 1.404E+07 | 19 | 398 | 417 | 240 | 1397 | 537 | 62% | 4022 nt (92.37) | 3.84x |
| | 2 | 7.779E+06 | 21 | 1555 | 1576 | 509 | | | | | |
| | 2* | 2.489E+07 | 17 | 5020 | 5037 | 651 | | | | | |
| HSJNC81 | 1 | 1.002E+07 | 136 | 4627 | 4763 | 2112 | 25536 | 1158 | 95% | 4044 nt (92.88) | 2.98x |
| | 2 | 1.632E+07 | 61 | 92177 | 92238 | 11152 | | | | | |
| | 2* | 3.147E+07 | 46 | 183602 | 183648 | 12273 | | | | | |
| HSJN240 | 1 | 1.550E+07 | 125 | 29996 | 30121 | 4239 | 7928 | 1692 | 79% | 4315 nt (99.1) | 15.36x |
| | 2** | 5.250E+06 | 170 | 18403 | 18573 | 3689 | | | | | |
| HSJN194 | 1 | 1.151E+07 | 10768 | NA | NA | 4381 | 4381 | 1215 | 72% | 2896 nt (89.9) | 30.8x |

**Supplementary Table 3. Mapping statistics for ancient viral genome reconstruction.** Viral enriched NGS libraries were sequenced and mapped to the human genome (hg19); unmapped reads are shown as total sequences. Number of reads kept in subsequent steps of B19V (COYC4, HSJNC81, HSJN240) and HBV (HSJN194) genome reconstruction are shown as well (described at Supplementary Methods 9). (*) A second independent round of capture was sequenced deeper in order to obtain a better coverage of the targeted viral ancient genome. (**) A different tooth was used to construct an independent library for the same individual. ($^{\#}$) Coverage and depth calculated based on the B19V CDS (AB550331) or whole HBV genome (GQ331046).

| Individual | Dental piece | Skeletal age (yr) | aDNA Genetic Sex | mtDNA haplogroup | Substrate for $^{14}C$ dating | $^{14}C$ Ages (BP +/- 1σ) CE |
|---|---|---|---|---|---|---|
| HSJNC81 | 1st Molar (Maxillary Left) | 24-34 | XY | L3d1b2 (African) | NA | NA |
| HSJN240 | 1st Premolar (Maxillary Right) | 35-50 | XY | L2a1b1a (African) | Third proximal phalange (left hand) | 408 +/- 30 |
| HSJN194 | 1st Molar (Maxillary Left) | 35-50 | XY | L0 (African) | Third proximal phalange (left hand) | 356 +/- 30 |
| COYC4 | 1st Molar (Maxillary Right) | NA | NA | B2 (Native-American) | NA | NA |

**Supplementary Table 4. Information for the human skeletal remains from which ancient viral genomes were recovered.**

a)

| Clock Model | Prior | Mean (95% HPD) Median root age in years | Median substitution rate (subs/site/year) | Likelihood | Posterior Probability |
|---|---|---|---|---|---|
| Strict | Exponential | 11282.97 (9952.19, 12684.59) | 1.04E-5 (8.76E-6, 1.20E-5) | -28102.68 (-28120.24, -28085.31) | -29342.09 (-29367.64, -29317.34) |
| | Constant | 11553.46 (10240.54, 13244.74) | 1.03E-5 (8.66E-6, 1-21E-5) | -28103.28 (-28122.22, -28086.40) | -29349.00 (-29375.42, -29323.36) |
| | Skyline | 11187.43 (9873.84, 12575.83) | 1.08E-5 (9.07E-6, 1.25E-5) | -28102.45 (-28120.45, -28085.73) | -29344.45 (-29372.69, -29318.07) |
| Relaxed-log | Exponential | 7913.17 (6980.99, 9748.39) | 1.96E-5 (1.19E-5, 2.81E-5) | -27744.36 (-27768.52, -27721.66) | -28926.25 (-28978.13, -28872.994) |
| | Constant | 9171.22 (7043.43, 13521.29)) | 2.62E-5 (1.50E-5, 3.98E-5) | -27743.42 (-27767.37, -27720.91) | -28894.34 (-28956.04, -28833.88) |
| | Skyline | 8824.31 (7040.62, 12155.6) | 2.04E-5 (1.45E-5, 2.71E-5) | -27743.89 (-27767.43, -27720.86) | -28916.19 (-28962.38, -28869.94) |

b)

| Clade | Median MRCA ybp (95% HPD interval) Strict, Exponential | Strict, Constant | Strict, Skyline | Relaxed-log, Exponential | Relaxed-log, Constant | Relaxed-log, Skyline |
|---|---|---|---|---|---|---|
| Genotype 1 | 7221.81 (70005.97, 7515.68) | 7245.25 (7012.39, 7551.84) | 7194.18 (6979.37, 7462.72) | 7230.44 (69929.16, 7807.76) | 7382.5 (6913.14, 8373.09) | 7356.44 (6940.73, 8163.19) |
| Genotype 2 | 2192.84 (1897.14, 2654.23) | 2219.73 (1891.33, 2700.34) | 2106.48 (1829.22, 2509.54) | 1878.53 (1473.94, 2504.52) | 1749.47 (1414.54, 2280.52) | 1720.87 (1438.26, 2142.25) |
| Genotype 3 | 3585.49 (3017.15, 4207.02) | 3642.21 (3045.81, 4333.63) | 3464.5 (2946.47, 4107.33) | 2350.11 (1412.72, 3526.24) | 2046.43 (1066.98, 3466.39) | 1932.92 (1205.45, 29899.18) |

**Supplementary Table 5. BEAST analysis of B19V evolutionary rates and MRCA.** Overview of B19V Bayesian evolutionary analyses with different molecular clock and priors used. **a)** comparison of median root age and substitution rates. **b)** comparison of MRCA per genotype.

| Sample | Lab code | Material | $^{87}Sr/^{86}Sr$ | 1 sd | 1 SE(M) | n | weight (g) | Sr Concentration (ppm) |
|---|---|---|---|---|---|---|---|---|
| Std EuA | SrT197 | standard | 0.708034 | 27 | 3 | 58 | | |
| 81A crown Lix1 | 6231 MA ID | enamel | 0.709419 | 37 | 5 | 52 | 0.00028 | 592.0 |
| 81A crown Lix2 | 6231 MA ID | enamel | 0.709508 | 27 | 4 | 57 | 0.00767 | 130.0 |
| 81A crown Res | 6231 MA ID | enamel | 0.710980 | 32 | 4 | 56 | 0.10709 | 127.3 |
| **81A parietal** | **6234 MA ID** | **bone** | **0.706718** | **32** | **4** | **58** | **0.11292** | **465.1** |
| 240 crown Lix1 | 6232 MA ID | enamel | 0.710920 | 41 | 6 | 47 | 0.00025 | 614.3 |
| 240 crown Lix2 | 6232 MA ID | enamel | 0.711024 | 29 | 4 | 56 | 0.00778 | 108.0 |
| 240 crown Res | 6232 MA ID | enamel | 0.711093 | 33 | 4 | 59 | 0.05398 | 173.0 |
| **240 phalange** | **6235 MA ID** | **bone** | **0.707553** | **36** | **5** | **57** | **0.03156** | **294.5** |

**Supplementary Table 6. $^{87}Sr/^{86}Sr$ in teeth (enamel) and bones in individuals HSJNC81 and HSJN240.** Lix1, Lix2 and Res correspond to the leaching stages. Errors during measurement are presented as one standard deviation with the last two digits (1 sd = $\pm 1\sigma_{abs}$). 1 SE(M) = 1sd/square root n. n = number of runs per analysis. Sr concentrations were determined with the Isotope Dilution technique. ppm = parts per million. The Eimer and Amend (EuA) Sr standard was analyzed during teeth and bone measurements. The certified $^{87}Sr/^{86}Sr$ value of this standard is 0.7080 ±0.0004 (Fairbairn et al., 1967).

| Group | Sample Nr | $^{87}Sr/^{86}Sr$ | Source | Location |
|---|---|---|---|---|
| A | WR [19564] | 0.70988 | Plutonic rock | Mali |
| | WR [19564] | 0.71204 | Plutonic rock | West African Craton (Mali) |
| B | WR [17605] | 0.7096 | Carbonatite | Algeria |
| | WR [17605] | 0.70972 | Carbonatite | Algeria |
| | WR [17605] | 0.70976 | Carbonatite | Algeria |
| | WR [17605] | 0.71007 | Carbonatite | Algeria |
| | WR [17605] | 0.71011 | Carbonatite | Algeria |
| | WR [17605] | 0.71125 | Carbonatite | Algeria |
| | WR [17605] | 0.71239 | Syenite | Algeria |
| C | WR [8695] | 0.70935 | Gabbro-diorite | West African Craton (Algeria) |
| | WR [8695] | 0.71016 | Gabbro | West African Craton (Algeria) |
| | WR [8695] | 0.71148 | Gabbro | West African Craton (Algeria) |
| D | J493a | 0.71183 | Granitic-dike | Ghana |
| | J509a | 0.70926 | Granite | Ghana |
| | KR8-066 | 0.71049 | Meta-greywacke | Ghana |
| | KR8-032 | 0.71163 | Meta-greywacke | Ghana |
| E | S007 | 0.71005 | Mafic intrusive | Senegal |
| | S005 | 0.71087 | Metavolcanic rock | Senegal |
| F | 12/449/01 | 0.70931 | Basalt | West African Craton |
| | 12/292/01 | 0.70957 | Dolerite | West African Craton |
| G | 3475 | 0.708395 | Soil | Mexico (Veracruz) |
| | 3476 | 0.708286 | Soil | Mexico (Veracruz) |
| | 3477 | 0.706688 | Soil | Mexico (Veracruz) |
| | 3478 | 0.706323 | Soil | Mexico (Veracruz) |
| | 3479 | 0.706614 | Soil | Mexico (Veracruz) |
| | 3480 | 0.705869 | Soil | Mexico (Veracruz) |

**Supplementary Table 7. $^{87}Sr/^{86}Sr$ Sources from West Africa and Trans Mexican Volcanic Belt (TMVB) used for calibration.** Compilation of $^{87}Sr/^{86}Sr$ from West Africa for groups A (Liégeois et al., 1991), B (Bernard-Griffiths et al.,

1988), C (Peucat et al., 2005), D (Ferrière et al., 2010), E (Fullgraf et al., 2013), and F (Sakyi et al., 2018); while G (Solís-Pichardo et al., 2017) correspond to the TMVB for isotopic comparison.

### Supplementary Figures:

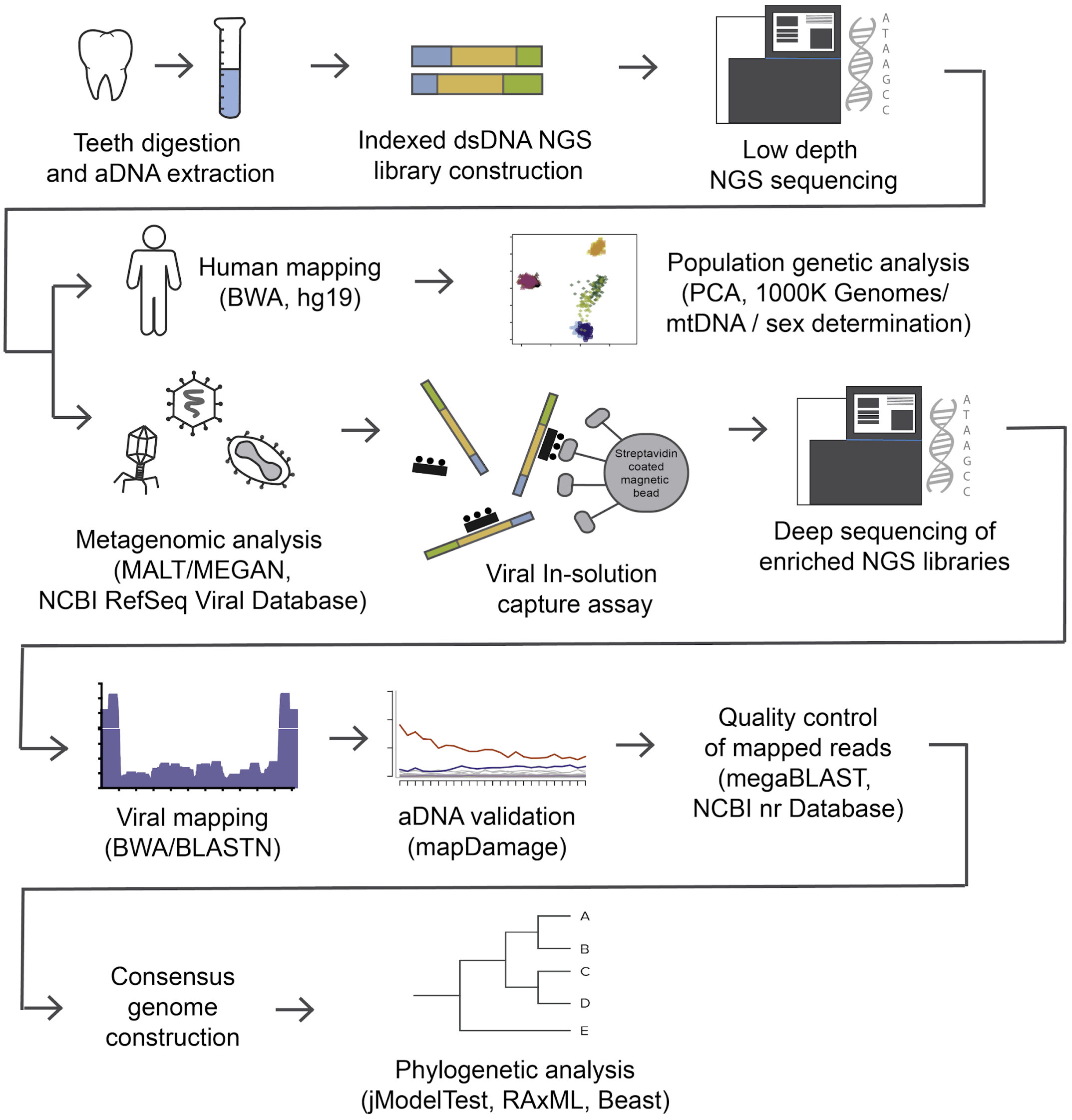


**Supplementary Figure 1. Pipeline followed for ancient viral genomes reconstruction.** Teeth were fragmented into small pieces to extract DNA from which NGS libraries were constructed. Reads were separated into human and non-human, for downstream analysis. After metagenomic analysis, samples were chosen for viral DNA enrichment and sequencing. Two independent viral DNA capture rounds were joined to reconstruct the ancient genome and estimate damage patterns. Consensus genome was constructed and aligned to a viral dataset with sequences from different genotypes and geographic regions to perform phylogenetic analysis.

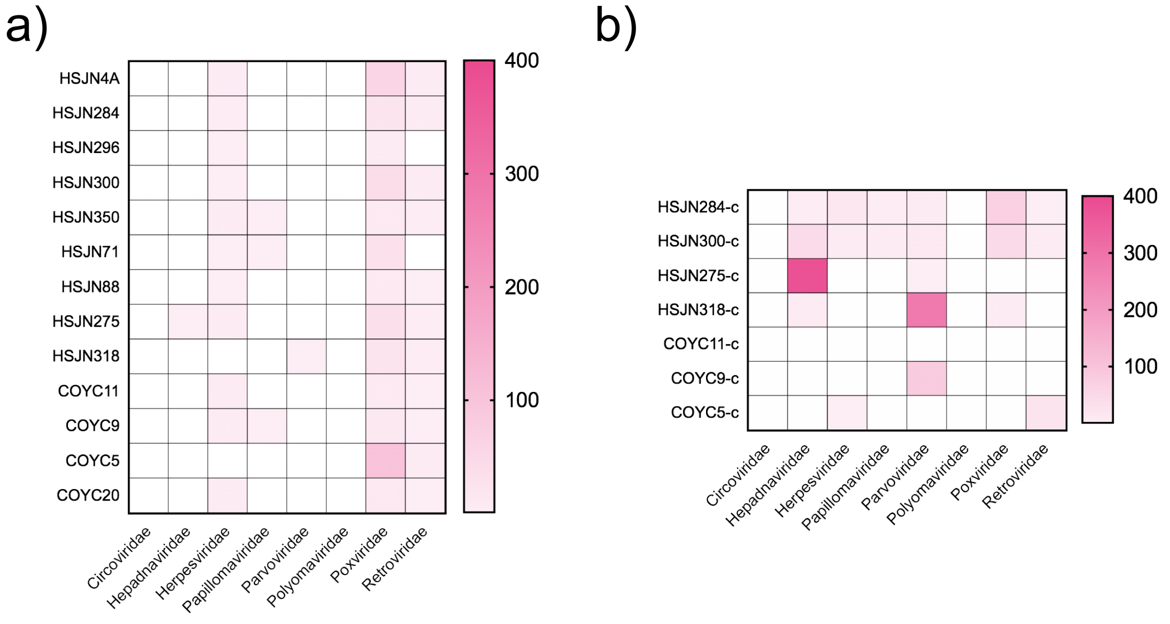


**Supplementary Figure 2. Individuals with DNA traces of clinical important viral families.** Metagenomic analysis performed on the Viral NCBI RefSeq Database with MALT, abundancies were normalized in MEGAN (default parameters). **a)** individuals with highest frequencies of hits to viral families included in our customized capture assay are shown, **b)** enriched samples normalized based on their respective shotgun sample in MEGAN (default parameters). Samples HSJN318 and HSJN275 were poorly enriched to cover the CDS region or present a clear damage pattern; respectively, hence not use for further phylogenetic analysis.

A)

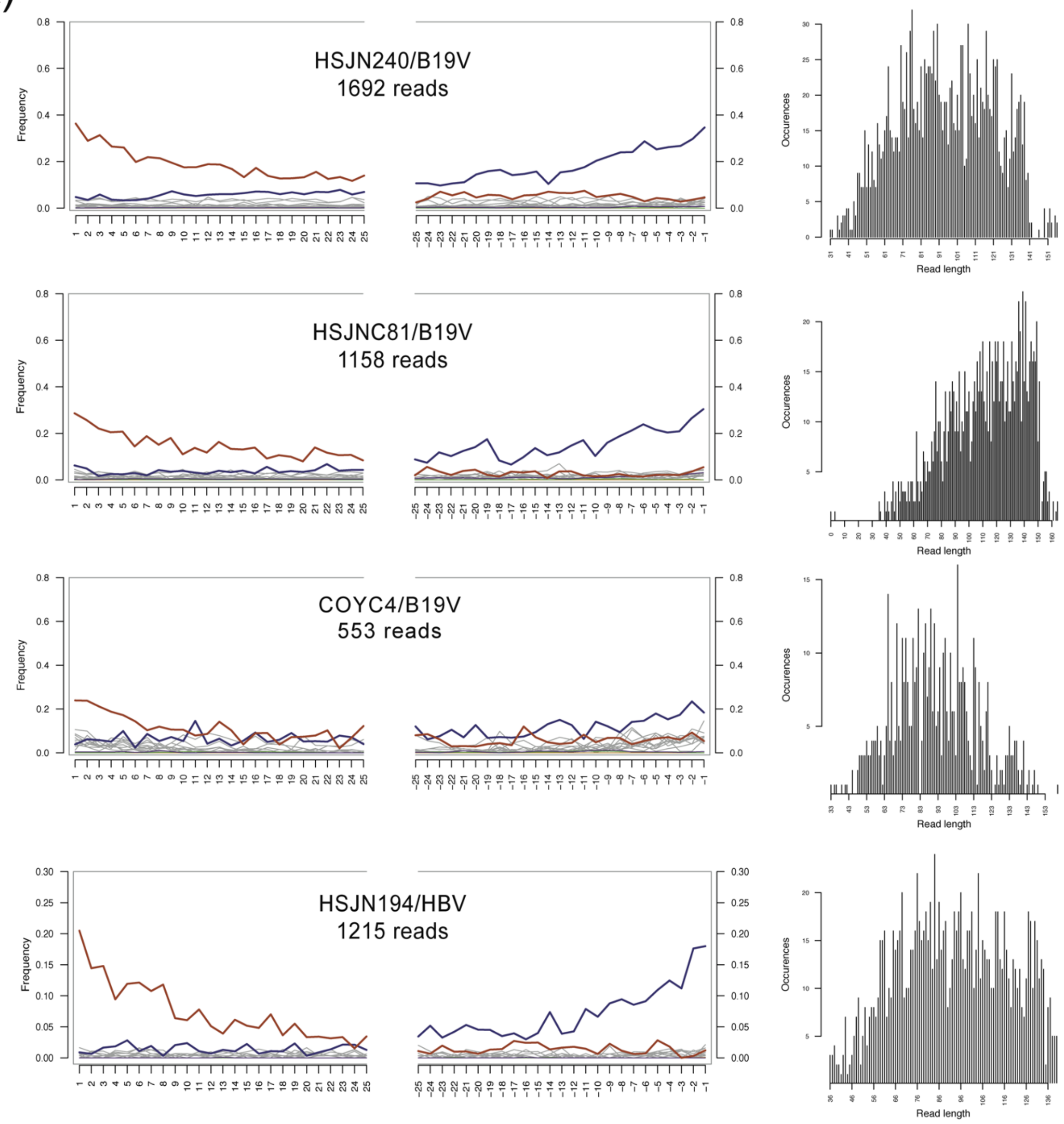
HSJN240/B19V
1692 reads
HSJNC81/B19V
1158 reads
COYC4/B19V
553 reads
HSJN194/HBV
1215 reads
Frequency
Occurences
Read length

B)

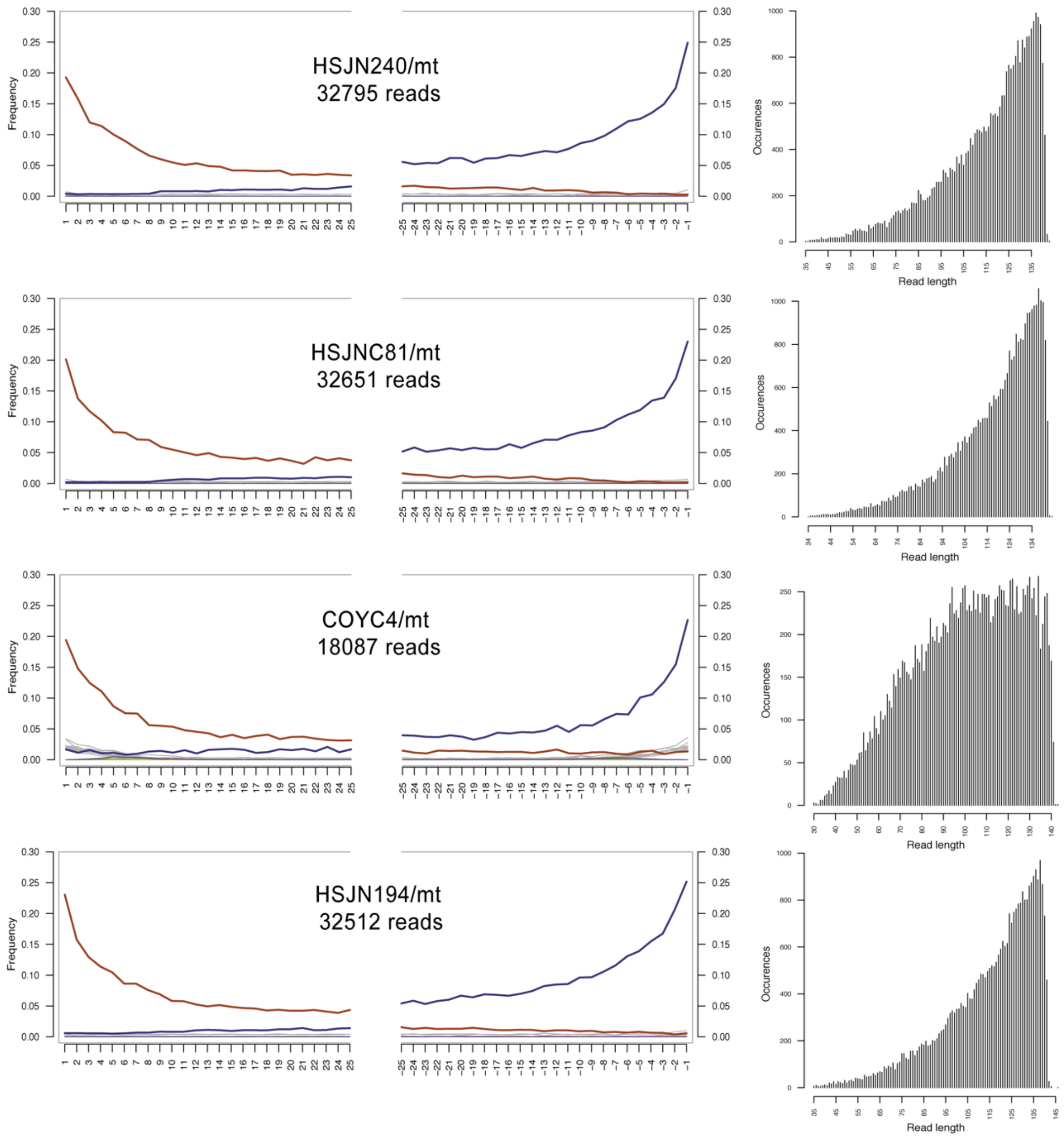


**Supplementary Figure 3. Damage patterns of ancient viral genomes and human hosts.** On the left; damage plots carried out with mapDamage 2.0; X axis shows the position (nt) on the 5’ and 3’ end of the read, on the Y axis the frequency of C>T and G>A transitions are indicated with red and blue lines, respectively while other mismatches are shown in gray. On the right, length distribution of the mapped reads without any cleavage treatment. Labels indicate individual/reference for a) virus (B19V or HBV); or c) mitochondria (rCRS). The number of mapped reads to the corresponding reference is indicated below each label.

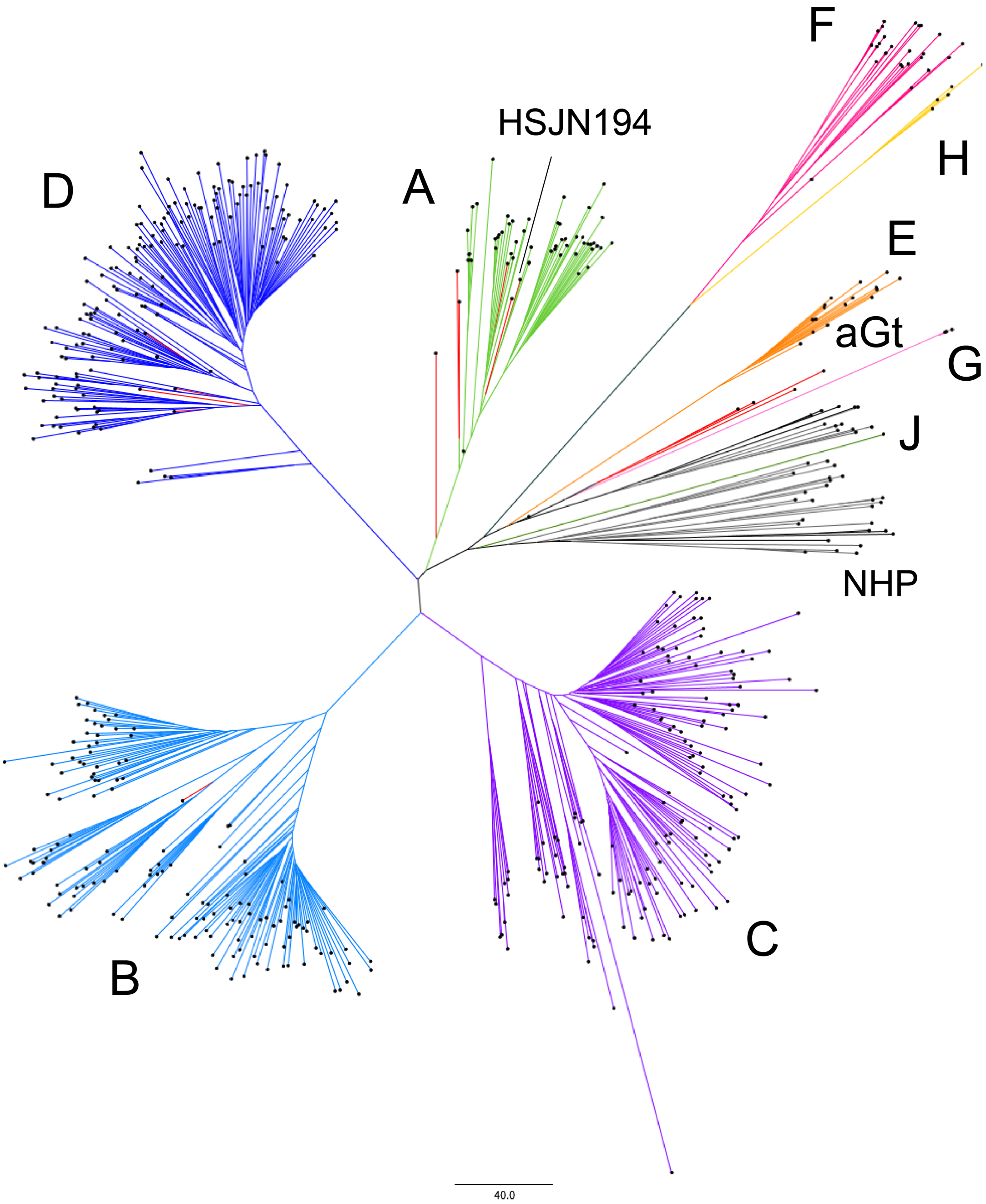


**Supplementary Figure 4. Phylogenetic analysis of HBV.** Modern sequences are colored by genotype; A: green; B: aqua; C: purple; D: blue; E: orange; F: magenta; G: pink; H: yellow; J: fern; Monkeys (primates not human): gray, while ancient genomes and aGt (ancient genotype) are in red. NHP: non-human primates. Neighbor joining tree constructed with MEGA based on a number of differences model with 1000 bootstraps with 604 genomes each represented with a black dot; tree was formatted on FigTree 1.4.3.

a)

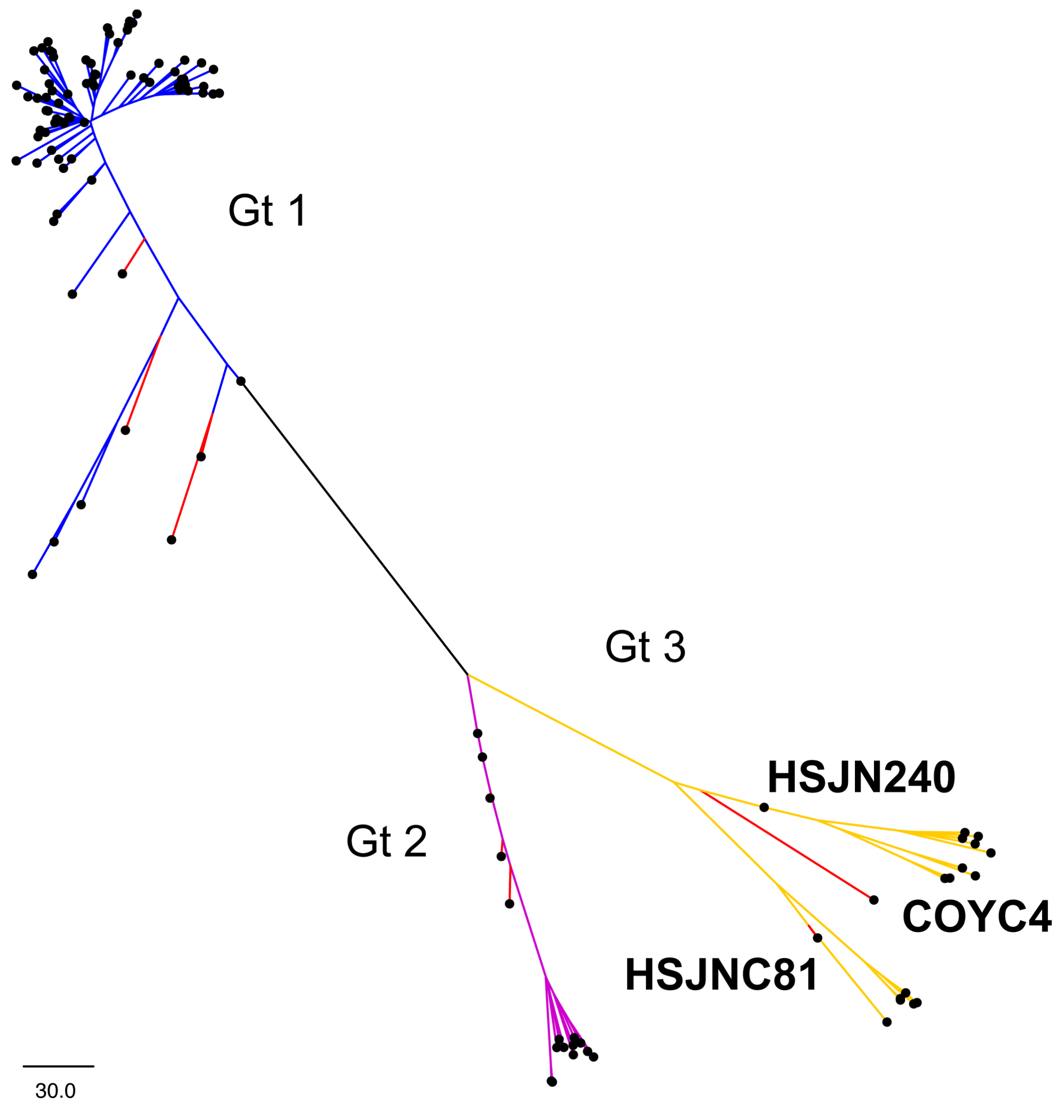

b)

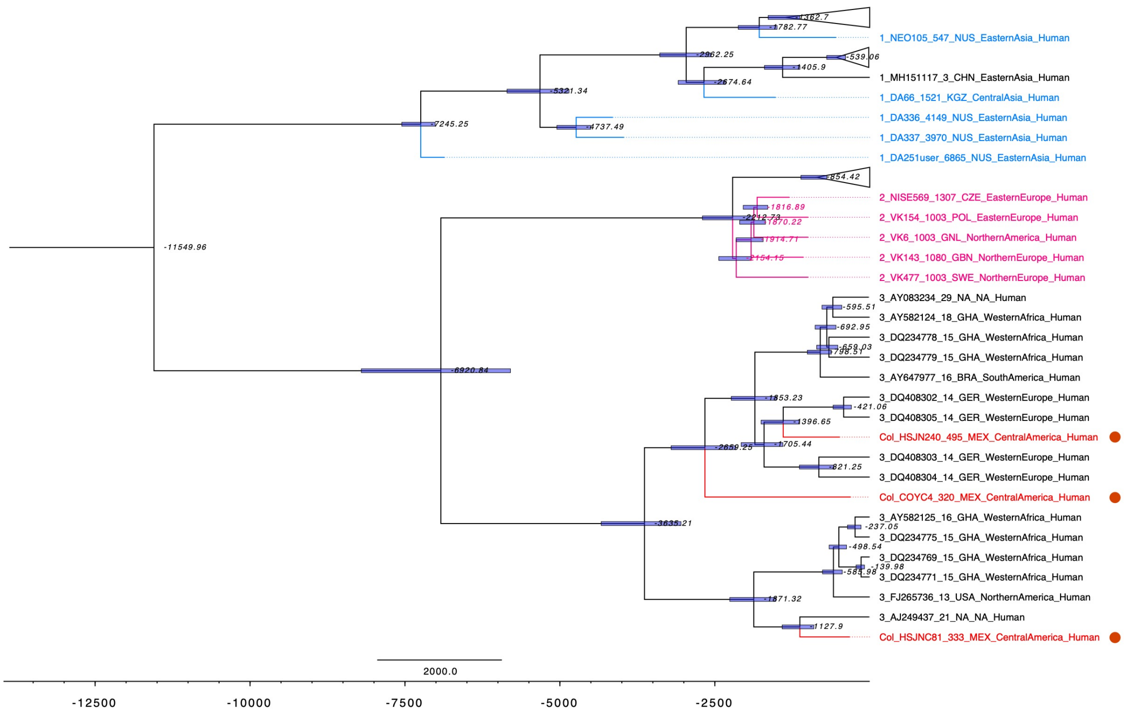


c)

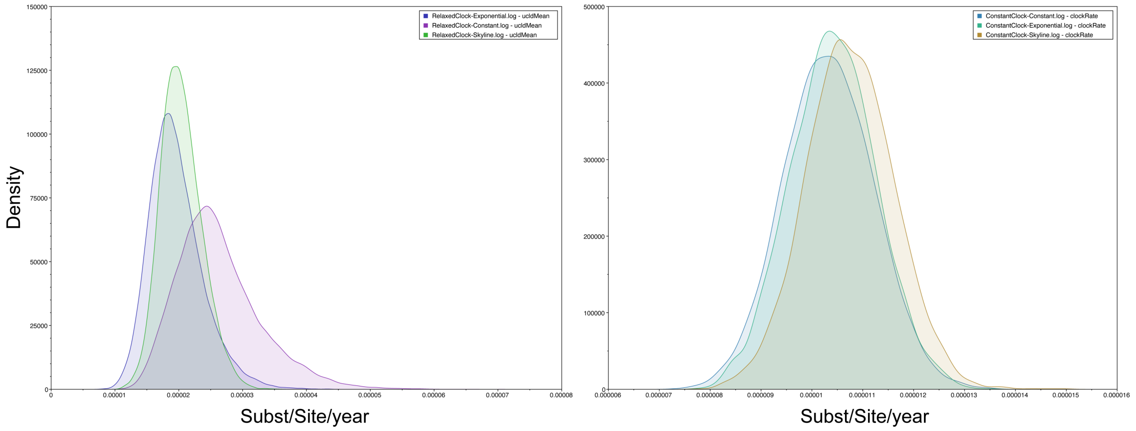


**Supplementary Figure 5. Phylogenetic analysis of B19V. a)** Neighbor joining tree constructed with MEGA based on a number of differences model with 1000 bootstraps. Genotypes 1, 2, 3 are shown with blue, magenta and yellow, respectively. Ancient genomes are shown in red; **b)** dated maximum clade credibility tree constructed with BEAST 2.5.1 using a constant population prior and a strict molecular clock; median node ages are shown in italics and 95% HPD shown as blue bars, modern genomes are black while ancient genomes are colored, colonial genomes from this study are shown in red, y-axis is shown in ybp, sequences from this study are highlighted with red circles. Both trees were formatted on FigTree 1.4.3. **c)** Posterior probability densities of mean evolutionary rate estimated with constant, exponential, Bayesian skyline population priors with a strict (left) and relaxed molecular clock (right).

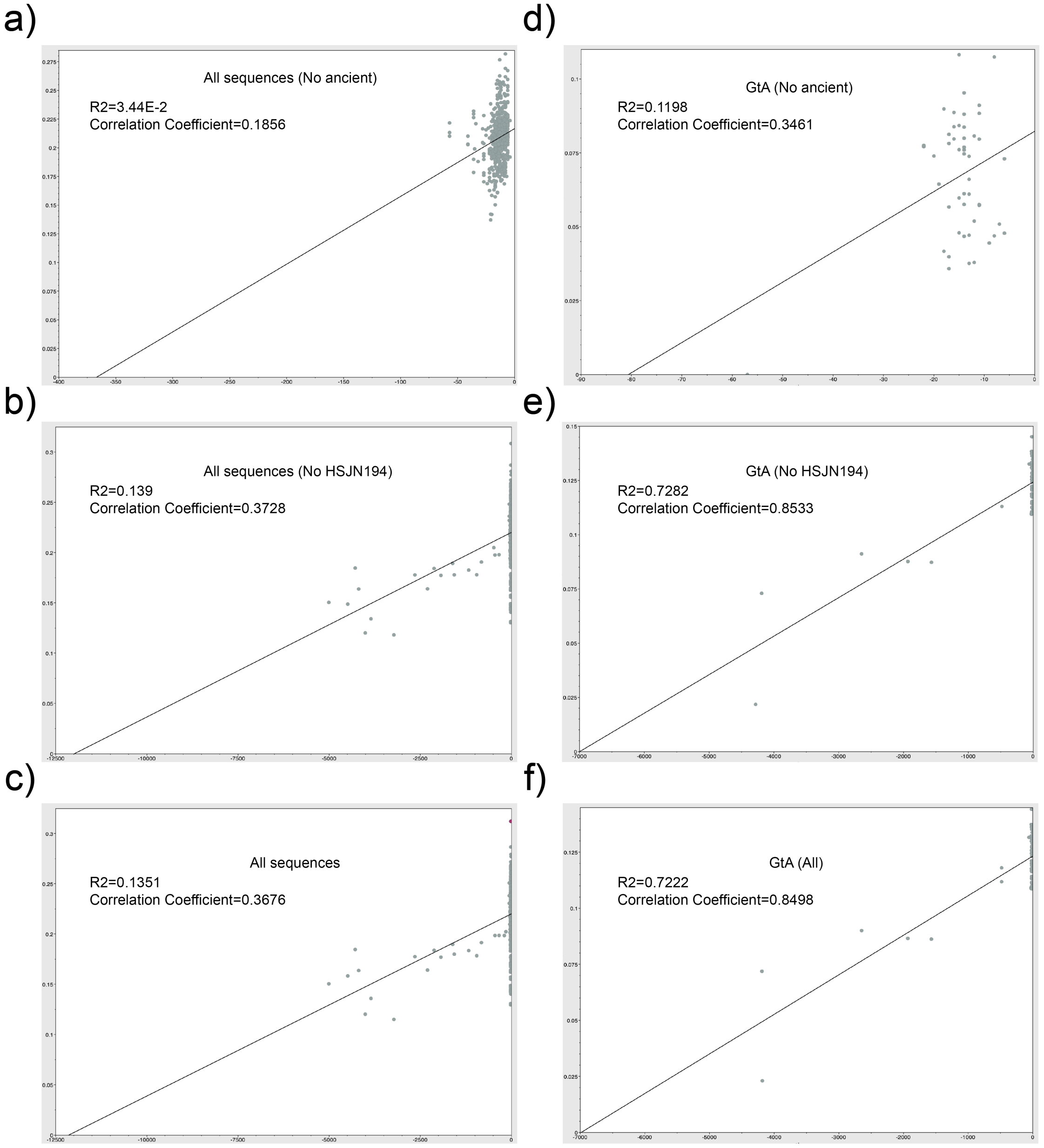


**Supplementary Figure 6. Root-to-tip regression analysis of HBV temporal structure.** Root-to-tip genetic distances on the y-axis were plotted against sampling time on the x-axis using Tempest 1.5.3. Each dot is an HBV genome in our ML analysis, and the central lines represents the regression line to which the evolution followed a clock-like pattern. Analysis were performed **a)** all sequences (HBV/DS2) without ancient **b)** without Colonial samples, and **c)** with all genomes (including ancient). Same approach was considered **d-f)** only for genotype A.

a)

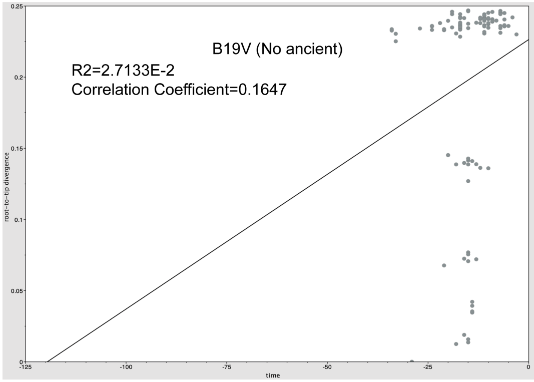


b)

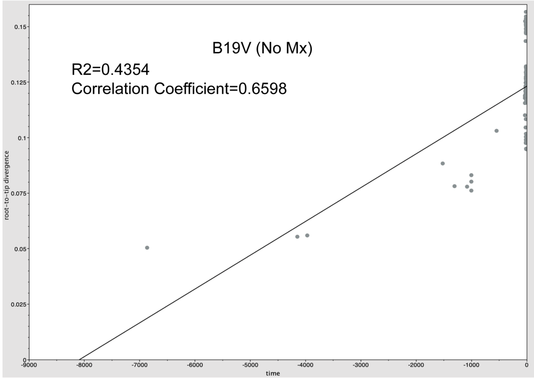


c)

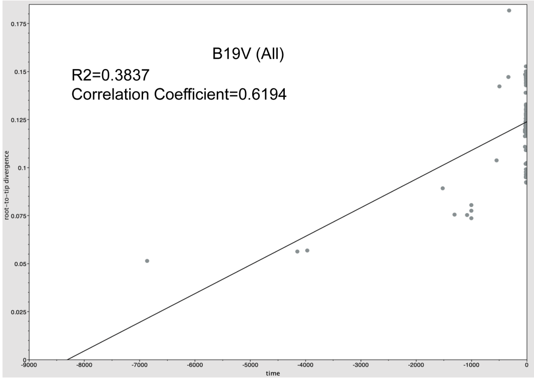


**Supplementary Figure 7. Root-to-tip regression analysis of B19V temporal structure.** Root-to-tip genetic distances were plotted against sampling time using Tempest 1.5.3. Each dot is an B19V genome in our ML analysis, and the central lines represents the regression line to which the evolution followed a clock-like pattern. Analyses were performed with B19V/DS2 including **a)** only modern sequences; **b)** without Colonial samples, and **c)** with all genomes (including ancient).

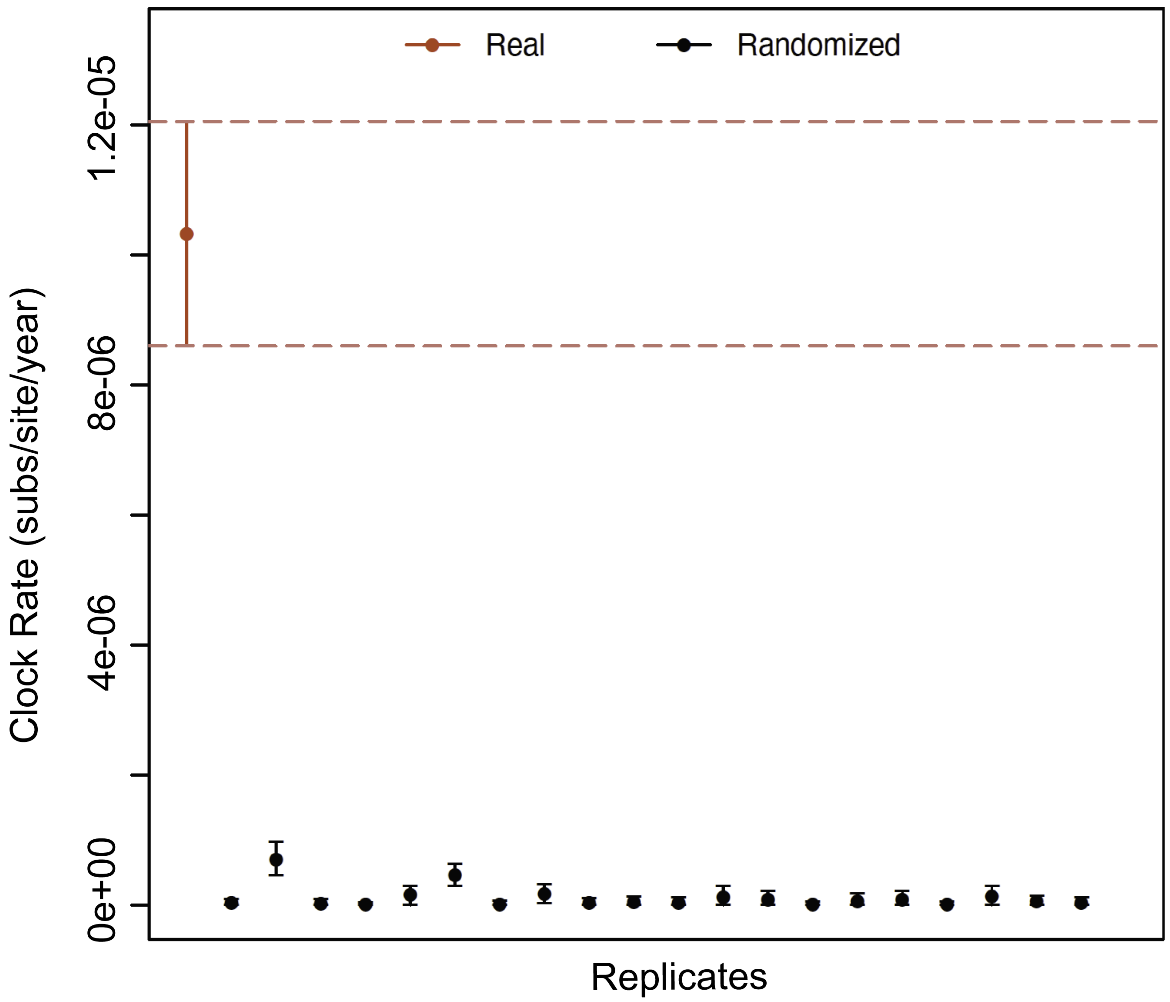


**Supplementary Figure 8. Date randomization test B19V.** Estimation of clock rates (subs/site/year) using 20 replicates with randomized dates (black dots) of the B19V/DS2 using a strict molecular clock and constant population prior. The dataset with the real sampling dates is shown as a red dot and the 95% credible intervals are shown as error bars.

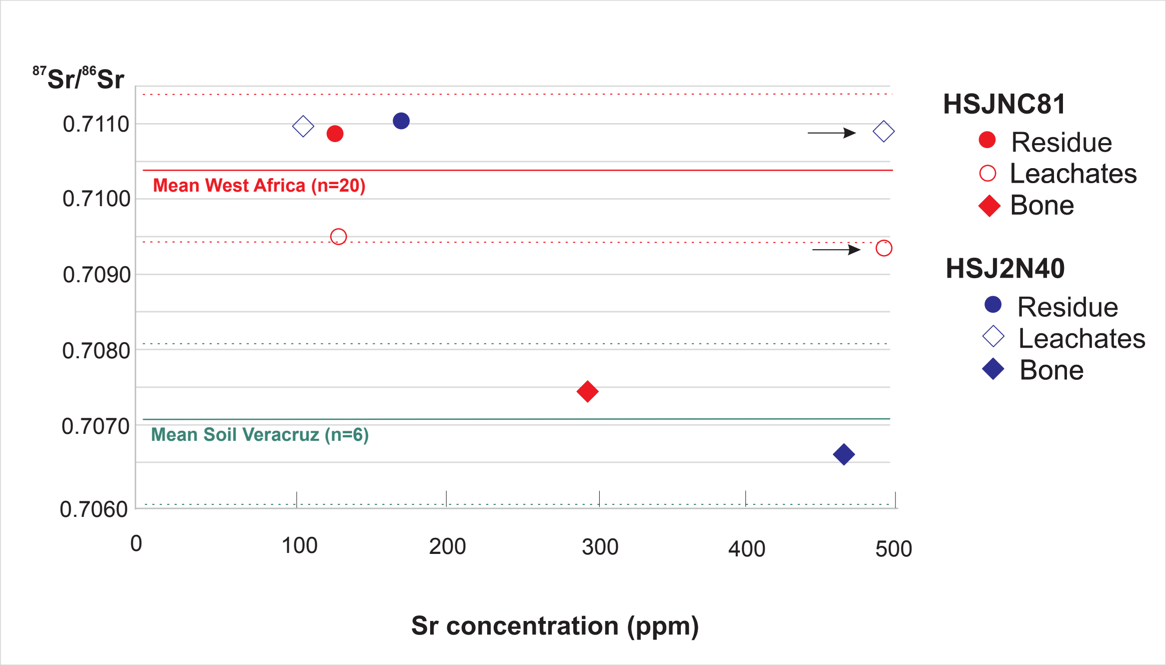


**Supplementary Figure 9. $^{87}Sr/^{86}Sr$ and Sr concentrations from HSJN individuals**. Analytical results are given in Supplementary Table 6. Arrowhead refers to second leachate values outside the graph. Red and green lines represent average $^{87}Sr/^{86}Sr$ values (solid lines) and 1 standard deviation (dashed lines) for West Africa igneous and metamorphic rocks (0.71041 ± 0.00099, Supplementary Table 7) and eastern TMVB soils in Veracruz, Mexico (0.70703 ±0.0011 (Solís-Pichardo et al., 2017)), respectively. The average $^{87}Sr/^{86}Sr$ of central TMVB soils (0.70449 ±0.00025, n=14 (Solís-Pichardo et al., 2017)) is not shown. Error bars are smaller than labels.

A)

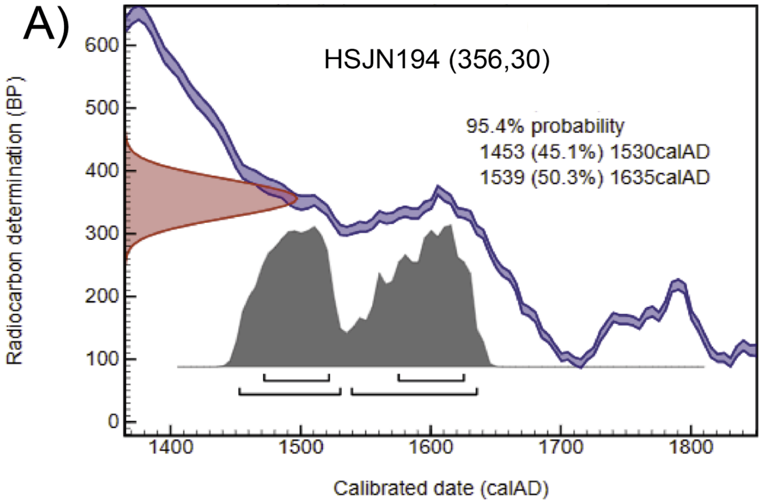


B)

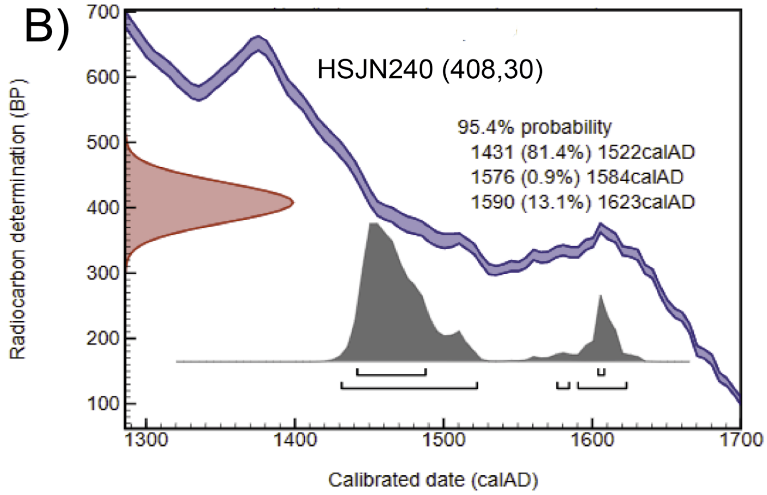


C)

| Sample | $^{14}$C Date Years (BP ± 1σ) | Calibrated date (Confidence Intervals) 1σ (68%) | 2σ (95%) |
|---|---|---|---|
| HSJN194 | 356 ± 30 | 479 – 325 cal BP<br>1472 - 1625 CE | 498 - 315 BP<br>1453 - 1635 CE |
| HSJN240 | 408 ± 30 | 509 – 343 cal BP<br>1442 - 1608 CE | 519 - 328 cal BP<br>1431 - 1623 CE |

**Supplementary Figure 10. Radiocarbon dating.** Radiocarbon dating calibrated on OxCal 4.3.2, based on IntCal13 atmospheric curve for HSJN194 (A) and HSJN240 (B). The radiocarbon determination of the samples is shown on the y axis, while the calibration based on known standards is shown as a double blue line. The calibrated date is on the x axis, and the likelihood age of the sample is shown as a gray solid distribution. A Summary of the calibrated interval dates is shown in (C). BP is considered as 1950 CE.

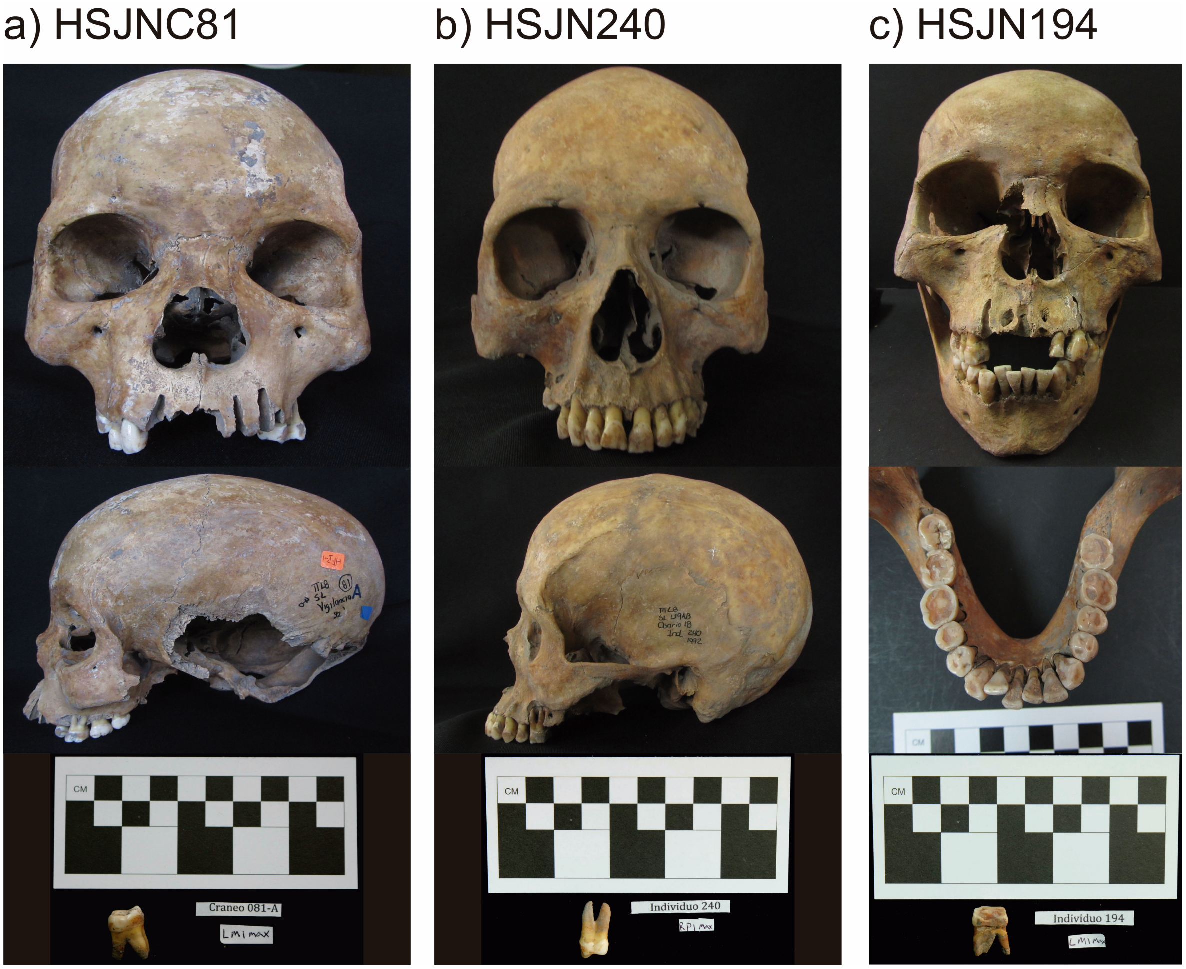


**Supplementary Figure 11. Individuals from the HSJN positive for ancient viruses.** Pictures from HSJN individuals and their respective teeth from which aDNA was extracted to reconstruct B19V (HSJNC81, HSJN240) and HBV (HSJN194) genomes. Only one tooth is shown for HSJN240.

Instituto Mexicano del Seguro Social.
